# Linguistic inputs must be syntactically parsable to fully engage the language network

**DOI:** 10.1101/2024.06.21.599332

**Authors:** Carina Kauf, Hee So Kim, Elizabeth J. Lee, Niharika Jhingan, Jingyuan Selena She, Maya Taliaferro, Edward Gibson, Evelina Fedorenko

**Affiliations:** Department of Brain and Cognitive Sciences, Massachusetts Institute of Technology, Cambridge, MA 02139 USA; McGovern Institute for Brain Research, Massachusetts Institute of Technology, Cambridge, MA 02139 USA; Department of Psychology, New York University, New York, NY 10012 USA; The Program in Speech and Hearing Bioscience and Technology, Harvard University, Cambridge, MA 02138 USA

## Abstract

Human language comprehension is remarkably robust to ill-formed inputs (e.g., word transpositions). This robustness has led some to argue that syntactic parsing is largely an illusion, and that incremental comprehension is more heuristic, shallow, and semantics-based than is often assumed. However, the available data are also consistent with the possibility that humans always perform rule-like symbolic parsing and simply deploy error correction mechanisms to reconstruct ill-formed inputs when needed. We put these hypotheses to a new stringent test by examining brain responses to a) stimuli that should pose a challenge for syntactic reconstruction but allow for complex meanings to be built within local contexts through associative/shallow processing (sentences presented in a backward word order), and b) grammatically well-formed but semantically implausible sentences that should impede semantics-based heuristic processing. Using a novel behavioral syntactic reconstruction paradigm, we demonstrate that backward- presented sentences indeed impede the recovery of grammatical structure during incremental comprehension. Critically, these backward-presented stimuli elicit a relatively low response in the language areas, as measured with fMRI. In contrast, semantically implausible but grammatically well-formed sentences elicit a response in the language areas similar in magnitude to naturalistic (plausible) sentences. In other words, the ability to build syntactic structures during incremental language processing is both necessary and sufficient to fully engage the language network. Taken together, these results provide strongest to date support for a generalized reliance of human language comprehension on syntactic parsing.

**Significance statement:** Whether language comprehension relies predominantly on structural (syntactic) cues or meaning-related (semantic) cues remains debated. We shed new light on this question by examining the language brain areas’ responses to stimuli where syntactic and semantic cues are pitted against each other, using fMRI. We find that the language areas respond weakly to stimuli that allow for local semantic composition but cannot be parsed syntactically—as confirmed in a novel behavioral paradigm—and they respond strongly to grammatical but semantically implausible sentences, like the famous ‘Colorless green ideas sleep furiously’ sentence. These findings challenge accounts of language processing that suggest that syntactic parsing can be foregone in favor of shallow semantic processing.

## 1. Introduction

Every day, humans produce and understand sentences they have never encountered before. This expressive power of language results from its compositionality: sentence meanings depend on the meanings of their constituent words and the ways in which the words relate to one another in the sentence’s structure. In particular, because sentence structure can be systematically decoded from the word order and/or morphosyntactic markers, we can understand novel inputs, even implausible/nonsensical ones, like the famous ‘*Colorless green ideas sleep furiously*’ example (Chomsky, 1957). However, the computations that enable us to quickly assemble complex meanings as we process language remain debated.

According to one view, humans perform ***rule-like symbolic parsing*** on linguistic inputs to derive complex meanings. Support for this view comes from strong sensitivity of human language processing mechanisms to structure. For example, a telltale signature of the language brain areas (Fedorenko et al., 2024) is a stronger response to structured stimuli, like sentences, compared to unstructured word-lists (Fedorenko et al., 2010; Pallier et al., 2011; Diachek et al., 2020; Shain, Kean et al., 2024), presumably because sentences engage computations related to structure building. This sensitivity to structure extends to even meaningless stimuli—so-called Jabberwocky sentences, like ‘*Twas brillig and the slithy toves did gyre and gimble in the wabe…*’ (Carroll, 1872)—compared to unstructured nonword-lists, although the overall response to nonword- composed stimuli is lower (Humphries et al., 2006; Fedorenko et al., 2010, 2016; Goucha & Friederici, 2015; Matchin et al., 2017; Shain, Kean et al., 2024). Another line of evidence comes from stronger responses in the language areas to more syntactically complex structures, including complexity associated with i) integrating words into a memory representation of the context (Holcomb, 1993; Ben-Shachar et al., 2003; Constable et al., 2004; Lau et al., 2006; Fedorenko et al., 2013; Blank et al., 2016; Shain et al., 2022; behavioral evidence: Gibson, 2000; Lohse et al., 2004; Grodner & Gibson, 2005) and ii) processing structurally unexpected elements (Lopopolo et al., 2017; Fedorenko et al., 2020; Shain, Blank et al., 2020; Heilbron et al., 2022; Goldstein et al., 2022; Wang et al., 2023; behavioral evidence: Demberg & Keller, 2008; Levy, 2008b; Smith & Levy, 2013; Shain et al., 2024).

An alternative family of views hold that human language comprehension may be more heuristic, shallow, and semantics-based than is often assumed (Ferreira et al., 2002; Sanford & Sturt, 2002; Tabor & Hutchins, 2004; Kim & Osterhout, 2005; Swets et al., 2008; Frank & Bod, 2011; Kuperberg, 2016; Mahowald et al., 2023). For example, Mollica, Siegelman et al. (2020) used fMRI to measure the language areas’ response to sentences with local word-order swaps (e.g., ‘*messages and gifts’è‘and messages gifts’*) and found that such manipulations do not decrease the response relative to well-formed sentences, even when sentences become quite syntactically degraded. Only when word swaps involved far-away words and destroyed local semantic dependencies, did the language areas’ response decrease. The authors therefore argue that interpretation can proceed without syntactic analysis, because our past linguistic experience tells us which words typically combine *semantically*. Thus, the fundamental computation of the language brain areas is ***syntax-independent semantic composition***.

However, Mollica, Siegelman et al.’s (2020) findings, and other evidence for syntax-independent semantic composition, have another explanation: humans may show robustness (and correspondingly strong responses in the language areas) to ill-formed inputs because they readily deploy error correction mechanisms (Gibson et al., 2013; Levy, 2008a, 2011; Levy et al., 2009; Zhang et al., 2024). Under this view, the human language system still performs syntax-driven composition, but syntactically corrupted input first needs to be reconstructed. Indeed, psycholinguistic studies have provided ample support for the human ability to cope with diverse errors, including word-order errors (Erickson & Mattson, 1981; Ferreira & Stacey, 2000; Ferreira & Bailey, 2004; Levy, 2008a; Mirault et al., 2018; Ryskin et al., 2018; Wen et al., 2021).

Here we test the necessity and sufficiency of syntactic processing for driving the brain’s response to language and find that the language network is fully engaged whenever syntactic structures can be built during incremental processing. These results argue against the shallower construals of linguistic computations.

## 2. Methods

Our approach is two-fold. First, we develop a novel behavioral paradigm that allows us to assess the ability to repair and interpret syntactically ill-formed input during incremental, word-by-word processing. Many past studies have examined the *costs* associated with the presence of syntactic violations using behavioral and neural measures (e.g., Osterhout & Holcomb, 1992; Hagoort et al., 1993; Newman et al., 2001; Kuperberg et al., 2003; De Vincenzi et al., 2003; Ditman et al., 2007; Friederici et al., 2010; Nieuwland et al., 2013). Other studies have investigated the *offline interpretation* of ill-formed linguistic inputs, where participants are presented with such inputs and have to answer questions that probe their interpretation of the sentence meaning or are asked to re-arrange the words into their most likely order (e.g., Ferreira & Stacey, 2000; Gibson et al., 2013; Chen et al., 2023, Mollica, Siegelman et al., 2020). However, to our knowledge, no method currently exists for evaluating how participants may be *interpreting ill-formed sequences as they process them incrementally*. We developed an approach where participants receive stimuli word by word (visually, on a computer screen) and, after each new word is uncovered, have the ability to reorder the words that are currently on display if they think the order in which they appear does not make sense (**Figure 1A**). By examining the orders that participants consider at different points in the sequence, we can make inferences about the syntactic structures they are building and the interpretations they are deriving. Using this new paradigm, we show that the stimuli with local word-order scrambling from Mollica, Siegelman et al.’s (2020) study, which elicit a strong response in the language brain areas, are amenable to real-time syntactic reconstruction, in contrast to conditions with more severe word-order rearrangement.

**Figure 1.**
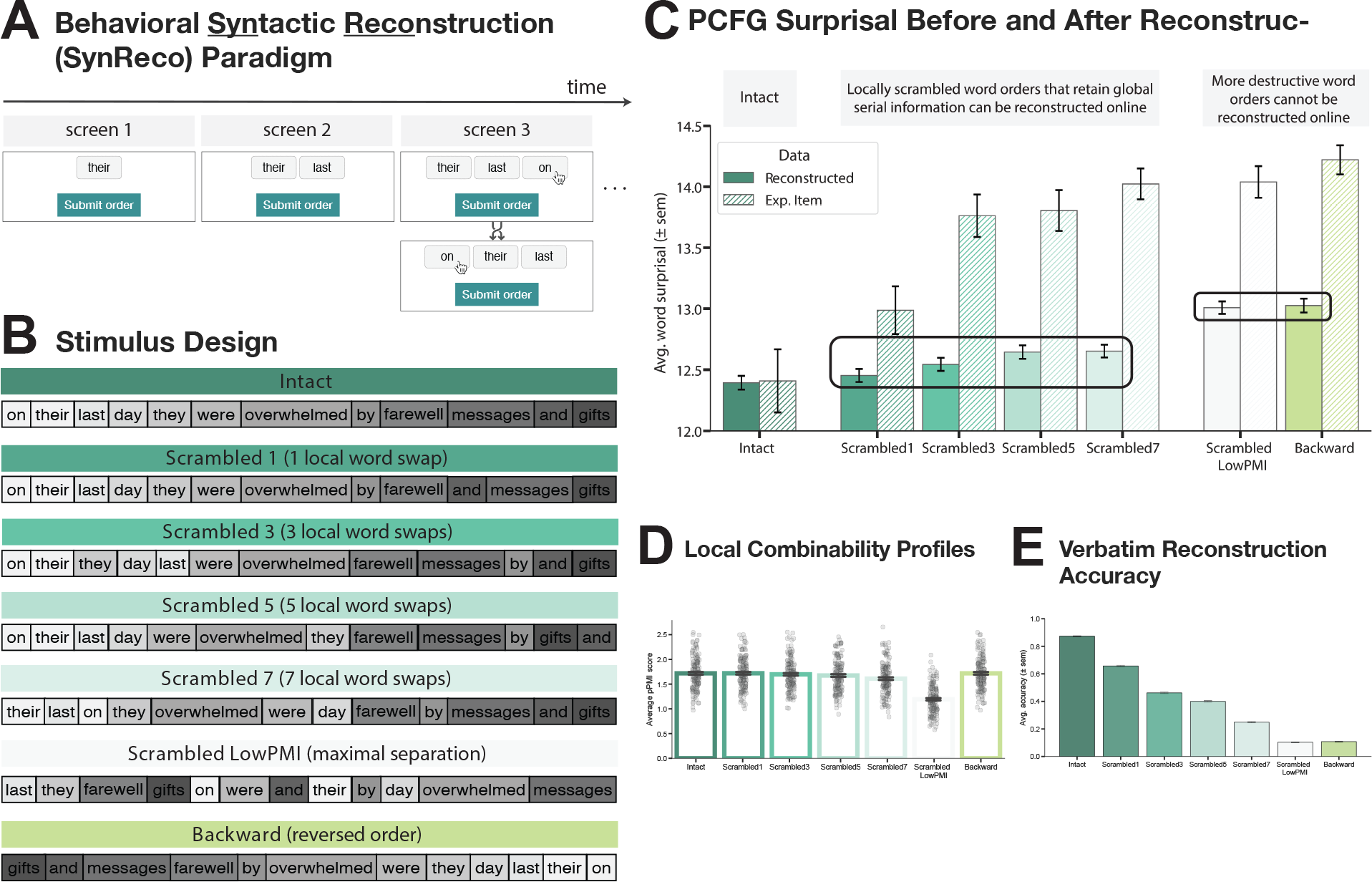
Word-order scrambled stimuli that elicit a sentence-level response in the language network are amenable to real-time syntactic reconstruction. A) Illustration of the novel behavioral <u>Syn</u>tactic <u>Reco</u>nstruction (SynReco) paradigm. Sentences are revealed on the screen one word at a time, in boxes with a drag-and-drop function. At each time step, participants can reorder the words on the screen into their best guess of the most likely word order. Each newly revealed word is appended to the participant’s last submitted order. **B)** A sample item from the critical experiment (figure adapted from Mollica, Siegelman et al., 2020); the grayscale color gradient is used to illustrate the increasing degradedness (i.e., the color spectrum becomes progressively more discontinuous with more swaps, but is preserved in the Backward condition). **C)** Average grammaticality of (i) the experimental materials (hatched bars) and (ii) participant reconstructions (solid bars) across conditions. Grammaticality is quantified as average PCFG word surprisal. Locally scrambled conditions are amenable to real-time reconstruction (PCFG surprisal following reconstruction is comparable to the *Intact* condition), but more severely degraded conditions (Scrambled *LowPMI* and *Backward*) are difficult to reconstruct in real time. **D)** Average positive PMI scores for experimental materials (see **Figure SI 4** for directional PPMI results). **E)** Average rate of recovery of the verbatim unscrambled input (reconstruction accuracy) after incremental parsing with the novel behavioral paradigm (verbatim reconstruction patterns replicate the results reported in Mollica, Siegelman et al., but as the fMRI results show, the PCFG surprisal estimates of the reconstructed strings (Figure 1C, solid bars), not the verbatim reconstruction accuracies, mirror the responses in the language network).

And second, using fMRI, we examine responses in the language areas to several linguistic conditions, including two critical conditions: (i) sentences presented backwards, from the last word to the first, and (ii) nonsensical but grammatically well-formed sentences. Sentences presented backwards allow for local semantic composition but should make syntactic reconstruction in real time challenging. In particular, reversing the words in a sentence does not disrupt local inter-word dependencies, so a processor that does not depend on word order should be able to perform local semantic composition. Consequently, the syntax-independent semantic composition hypothesis predicts that the response in the language areas should be as high as to grammatical sentences. However, the consistent reversal of the order of constituent elements intuitively places a substantial burden on the parsing mechanisms, at least in a language that relies heavily on word-order cues (as confirmed empirically, using our novel behavioral paradigm described above). Therefore, the syntax-dependent composition hypothesis predicts a low response in the language areas.

In contrast, nonsensical but grammatically well-formed sentences (like the ‘*Colorless green ideas…*’ example; Chomsky, 1957) allow for syntactic structure building, but building complex meanings is impeded by semantic implausibility. As noted in the <u>Introduction</u>, the response in the language areas to typical (well-formed and plausible) sentences is higher than to Jabberwocky sentences, which are syntactically well-formed but made up of nonwords (Humphries et al., 2005; Fedorenko et al., 2010; Shain, Kean et al., 2024). But is the stronger response to typical sentences due to i) the presence of real words, which are the stored basic units of language and contain critical cues for structure building (e.g., Pollard & Sag, 1994; Steedman, 2000; Goucha & Friederici, 2015), or ii) the plausibility of the resulting meanings, which can aid top-down semantic prediction (e.g., Kutas & Hillyard, 1984; Federmeier & Kutas, 1999; Nieuwland & Van Berkum, 2006; Bicknell et al., 2010; Matsuki et al., 2011)? The syntax-independent semantic composition hypothesis, whereby interpretation proceeds ‘bottom up’ from local semantic associations, predicts a low response to nonsensical sentences in the language areas because our **language processor should not attempt to form complex meanings from words that are unlikely to enter into a semantic dependency** (based on our prior linguistic experience and world knowledge). In contrast, the syntax-dependent semantic composition hypothesis predicts a strong response given that semantic composition can still take place even if the resulting meaning is implausible. If it turns out that the language areas’ computations are not affected by plausibility, this would open up new questions about where in the brain plausibility is computed.

### 2.1 Experiment 1: Behavioral incremental syntax reconstruction study

In this experiment, we assessed people’s ability to reconstruct syntactically ill-formed input during incremental word-by-word processing.

#### 2.1.1 Design and materials

We developed a novel behavioral paradigm (SynReco for “<u>syn</u>tax <u>reco</u>nstruction”) in which sentences are revealed one word at a time, with each word presented within a box that has a drag-and-drop function. At each time step, participants are asked to reorder the words on the screen according to their best guess as to the most likely word order, even when the current displayed sentence fragment does not yet form a grammatical sequence. Participants can reorder as many words as they want, until they are satisfied with the resulting order. Once they are satisfied with the order, they can click a “Submit order” button to reveal the next word or, if the last word of the stimulus has already been revealed, to end the trial.

The experiment included intact sentences and different versions of altered-word-order sentences. The critical stimuli consisted of 35 items, each in 7 versions (corresponding to conditions), for a total of 245 stimuli. The stimuli were adopted from Mollica, Siegelman et al. (2020). As described in greater detail in Mollica, Siegelman et al. (2020), a set of 12-word-long sentences (the **Intact** condition) were extracted from the British National Corpus (BNC; Leech, 1992), of which we used a subset of 35 sentences. For consistency with the materials used in our fMRI study (see Section 2.2.1), we converted the word spellings to American English (e.g., *moustache* è *mustache*). Five of the six altered-word-order conditions came directly from Mollica, Siegelman et al. (2020): The locally scrambled conditions, **Scrambled{1,3,5,7}**, were created by iteratively and randomly choosing 1, 3, 5, or 7 words in each original sentence and swapping them with one of their immediate word neighbors. As reported in Mollica, Siegelman et al. (2020), these local word-swap manipulations, even for the 7-swap case, typically preserve local semantic dependency structure, as can be measured by pointwise mutual information (PMI) among nearby words (as detailed in Section 2.2.1). The **Scrambled_LowPMI** condition minimizes the combinability among nearby words and was created by placing content words that are adjacent/proximal in the original sentence far away from each other. In addition to these six conditions, we included a novel (**Backward**) condition, which was created by reversing the word order in the original sentences. *Backward* sentences are characterized by the same local semantic dependency structure as their *Intact* counterparts (**Figure 1D**, **Figure SI 1**, **Table SI 1**). The 245 stimuli were distributed across 7 experimental lists (35 stimuli each, 5 per condition) such that each list contained only one condition of an item. Each participant completed only one list.

Each list additionally included 7 practice items, and 19 filler items (7 of which were designed to be easier than the critical stimuli and served as attention checks). These items differed from the critical items in length (mean = 7 words, SD = 1.5), but, similar to the critical items, they were either grammatically well-formed or contained one or several local word swaps. We varied the number of words in these non-critical items in order to prevent participants from relying on their expectation of how many words are still to come in a sequence while performing the critical word- reordering task.

#### 2.1.2 Procedure

The experiment was implemented as a new JQuery module within the Ibex web-based psycholinguistic experiment software platform (https://github.com/addrummond/ibex).

The experiment began with detailed instructions. To encourage the careful reading of these instructions, we divided the information across three screens and allowed participants to advance to the next screen only after a 15-second delay, when a “Continue” button appeared. Participants were told that their task is to try to create a grammatical word order at each point during the trial. They were encouraged to try their best before moving on to reveal the next word, but they were also told that if—at some point in the trial—they cannot find a way to reorder the words so as to form a grammatical string, they should reveal the next word to see if that helps. For example, if the first two words are “this a”, reordering them does not help (“a this” is not a grammatical string), but the next word may be “is”, in which case a grammatical string can now be formed (“this is a”). Participants were informed that the words in each trial could always be reordered into (at least one) fully grammatical sentence (i.e., the original version of the sentence) and were advised that (i) on some trials no reordering may be needed, and that (ii) some trials might prove challenging. They were also told that in some cases, there may be multiple ways to order the words so as to create a grammatical string (e.g., “books and pencils” vs. “pencils and books”), and that in such cases, any of the permissible word orders would be accepted. Finally, they were warned that the experiment includes several (unmarked) attention check items—used to assess and maintain participants’ attentiveness throughout the experiment—and that consistent failure on these items might lead to exclusion from the experiment, whereas consistently good performance throughout the experiment would lead to a small bonus payment.

Following the instructions, participants completed 7 practice items. All stimuli were presented in lowercase letters and without punctuation. For the practice items, i) participants received feedback at each time step telling them if the submitted word order was permissible or not, and ii) the next word in the sequence was only revealed once participants had submitted one of the permissible word orders for the current time step. To be able to do this, we had to determine all grammatically allowed word orders at each time step for each of these items (as well as for the filler items—see below). C.K. and J.S. made an initial set of judgments about the word orders, and these judgements were then validated in an independent experiment (n = 23 participants). In particular, participants completed the experiment in the same experimental procedure as the critical experiment, and J.S. manually reviewed the set of unique orders submitted by the participants at each time step to include any additional grammatically allowed word orders that were missing from the initial set of permissible orders. In this way, the practice items served to train participants to submit their guess of the most likely word order *at each time step*, even if that guess may turn out to be wrong later on, instead of revealing multiple words at once before starting to reorder them.

Upon the completion of the practice items, the critical experiment began. For the critical items, participants did not receive any feedback; for the filler items, which were randomly interspersed with the critical items, they received feedback. In particular, for the subset of the filler items that served as attention checks, participants were notified when they submitted an ungrammatical word order at any point in the trial, to encourage them to carefully arrange the words at each time step. If they submitted grammatical orders at each time step, they were notified at the end of the trial that they passed an attention check. For the remaining filler items, participants were informed of their performance (pass or fail) when they submitted the final word order. The average completion time for this experiment was ∼50 min.

#### 2.1.3 Participants

We recruited 140 participants through the Prolific web-based testing platform, restricting our task to participants with IP addresses in the United States. Participants were included in the analyses if they satisfied all of the following criteria: (i) they succeeded on at least 4 of the 7 attention check items (see above for details), and (ii) they succeeded on at least 6 of the 12 remaining filler items (see above for details). The numbers for i and ii were determined based on the error distributions (excluding the lowest quartile of participants, see **Figure SI 2**). Data from 72 participants were included in the final analysis.

### 2.2 Experiment 2: fMRI study

Each participant completed (i) the critical task, (ii) a language network localizer task, which was used to identify language-responsive brain regions (Fedorenko et al., 2010), and (iii) a localizer task for another network: the domain-general Multiple Demand (MD) network (Duncan, 2010), which was used in some control analyses. Most participants also completed one of two tasks for unrelated studies. The scanning sessions lasted approximately two hours.

#### 2.2.1 Critical task design and materials

Participants passively read 12-word-long stimuli in a blocked design. The stimuli were presented one word/nonword at a time and belonged to one of eight conditions: (1) Intact plausible sentences (S), (2) Backward sentences (BS), (3) Nonsense sentences (NS), (4) Jabberwocky sentences (JS), (5) Word lists (WL), (6) Nonword lists (NWL), and two conditions used to address a distinct research question ((7-8) Predictable and unpredictable phrase lists; see <u>Discussion</u>).

The stimuli for the sentence conditions consisted of 192 items, each in 4 versions (corresponding to conditions: S, BS, NS, and JS), for a total of 768 stimuli. The base set (for the **Intact plausible sentences (S)** condition) consisted of 140 sentences adopted from Mollica, Siegelman et al. (2020) and an additional set of 52 sentences. As described in greater detail in Mollica, Siegelman et al. (2020), a set of 12-word-long sentences were extracted from the British National Corpus (BNC; Leech, 1992), of which we used a subset of 140 sentences (the remaining 10 sentences in the Mollica, Siegelman et al.’s study contained a high proportion of function words, which presented a challenge in creating satisfactory *Nonsense* sentence variants; see below for details). We additionally extracted a set of 52 12-word-long sentences from the same corpus. We made minor adjustments to some of the sentences by converting the word spellings to American English (e.g., *moustache* è *mustache*).

To create the **Backward sentences (BS)** condition, we reversed the order of the words in each of the 192 sentences. A critical design feature of the stimuli in this condition is that they are characterized by the same local syntactic and semantic combinability of nearby words as the original sentences (see **Figure 1D; Figure SI 2**).

To create the **Nonsense sentences (NS)** condition, we first created a set of candidate nonsense sentences, where for each of the 192 sentences, we replaced each content word (noun, verb, adjective, and adverb) with a replacement word. Suitable replacement words were determined based on morpho-syntactic feature overlap with the word to be replaced. Specifically, we classified each word in each sentence according to its part-of-speech tag, syntactic dependency label, and morphological features (such as case, number, mood, or tense information), all obtained using the NLTK (Loper & Bird, 2002) and spaCy (Honnibal et al., 2020) models. Verbs were further subcategorized according to their argument valence, to approximate the linguistic contexts in which they could serve as possible replacements for other verbs. Nouns were additionally annotated with coarse phonological features, such as whether they began with a vowel or a consonant sound (determined via their orthography as a proxy) to ensure that the replacement nouns would have the right onset when following indefinite determiners. Based on these annotations, we created a “dictionary” that mapped a given feature set to a list of all content word tokens (with duplicates; e.g., if a word “cat” occurred twice in the original set of sentences, it was included twice in the dictionary) and consisted of 1,398 content words. We then iterated through the sentences and replaced each content word with a randomly sampled word from our dictionary that corresponded to the original word in terms of its annotated features. In this way, the component words were the same between the S and the NS conditions. The algorithm was largely successful based on examining the resulting sentences; however, we carefully hand-checked the algorithmically-created nonsense sentences to ensure that the category selectivity of the verbs was satisfied, because the unique distributional signature of each verb could only be coarsely approximated by the feature annotations. Hence, if needed, we chose another replacement word from the dictionary.

To create the **Jabberwocky sentences (JS)** condition, for each of the 192 sentences, we replaced each content word with a suitable replacement nonword. Suitable replacement nonwords were generated using the *generate classic* method for English from the Wuggy pseudoword generator python package (Keuleers & Brysbaert, 2010). This algorithm creates nonwords by matching the original word in terms of sub-syllabic structure and syllable-transition frequencies. The latter constraint ensures the preservation of functional morphology (e.g., the past tense marker -ed or the plural -s), because replacing these high-frequency syllables typically involves a massive change in transition frequency (Keuleers & Brysbaert, 2010). We then iterated through the sentences and replaced each content word with a matching nonword.

To create the stimuli for the **Word lists (WL)** condition, we gathered all words across the 192 sentences and randomly recombined these 2,304 words (192 sentences, each 12 words long) into 192 sequences of 12 words each (via sampling without replacement). To create the stimuli for the **Nonword lists (NWL)** condition, we gathered all words and nonwords across the 192 Jabberwocky sentences and randomly recombined these 2,304 words/nonwords into 192 sequences of 12 words/nonwords each (via sampling without replacement). The stimuli for the remaining two conditions, which were used to address a distinct question (see <u>Discussion</u>), consisted of 12-word-long sequences, each made up of six determiner-noun phrases of the form ‘the noun’ (see **Figure SI 13** for details).

The stimuli for the sentence conditions (S, BS, NS, JS; 192 stimuli per condition)—where correspondence exists among the different-condition versions of the same sentence—were distributed across four experimental lists, such that each list contained only one version (S, BS, NS, or JS) of a given sentence and 48 stimuli for each of the four sentence conditions. In addition to the 192 sentence stimuli, each list included 48 Word Lists (WL), 48 Nonword Lists (NWL), and 96 items across two conditions that are not relevant to the current study (see <u>Discussion)</u>, for a total of 384 stimuli. Each participant completed only one list.

Prior to the experiment, we ensured that stimuli across conditions did not differ in low-level features, such as word frequencies or word lengths (see **Figure SI 3**). We additionally evaluated the stimuli to ensure that they have the desired properties for dissociating the syntax-dependent and the syntax-independent semantic composition hypotheses, with a focus on the two critical conditions (see **Figure 2C**): *Backward* sentences and *Nonsense* sentences (the original sentences are used for comparison). First, we examined two **model-derived** measures to determine the degree to which a string supports *syntax-driven sentence-structure building*. We used probability estimates from (i) a lexicalized, probabilistic context-free grammar (PCFG) model (Booth, 1969), and (ii) a powerful neural language model, GPT2-xl (Radford et al., 2019). These two measures are complementary: whereas the PCFG model computes probability estimates based on the structured syntactic representation while ignoring surface-level patterns of word co-occurrence, the GPT2-xl model does the reverse: it computes probability estimates solely based on the surface-level patterns of word co-occurrence, and only implicitly considers information about sentence structure. To derive lexicalized **PCFG surprisal** scores, we use the incremental left-corner parser of Van Schijndel & Schuler (2013), trained on a generalized categorial grammar (Nguyen et al., 2012) reannotation of Wall Street Journal sections 2 through 21 of the Penn Treebank (Marcus et al., 1993) (see also Shain, Blank et al., 2020). To derive a single score per sentence, we obtained a surprisal estimate for each word, and then averaged these estimates across all words in the sentence. The surprisal scores under the **GPT2-xl** model were calculated as the average token surprisal derived from the pre-trained model checkpoint available through the HuggingFace transformers library (Wolf et al., 2020).

**Figure 2.**
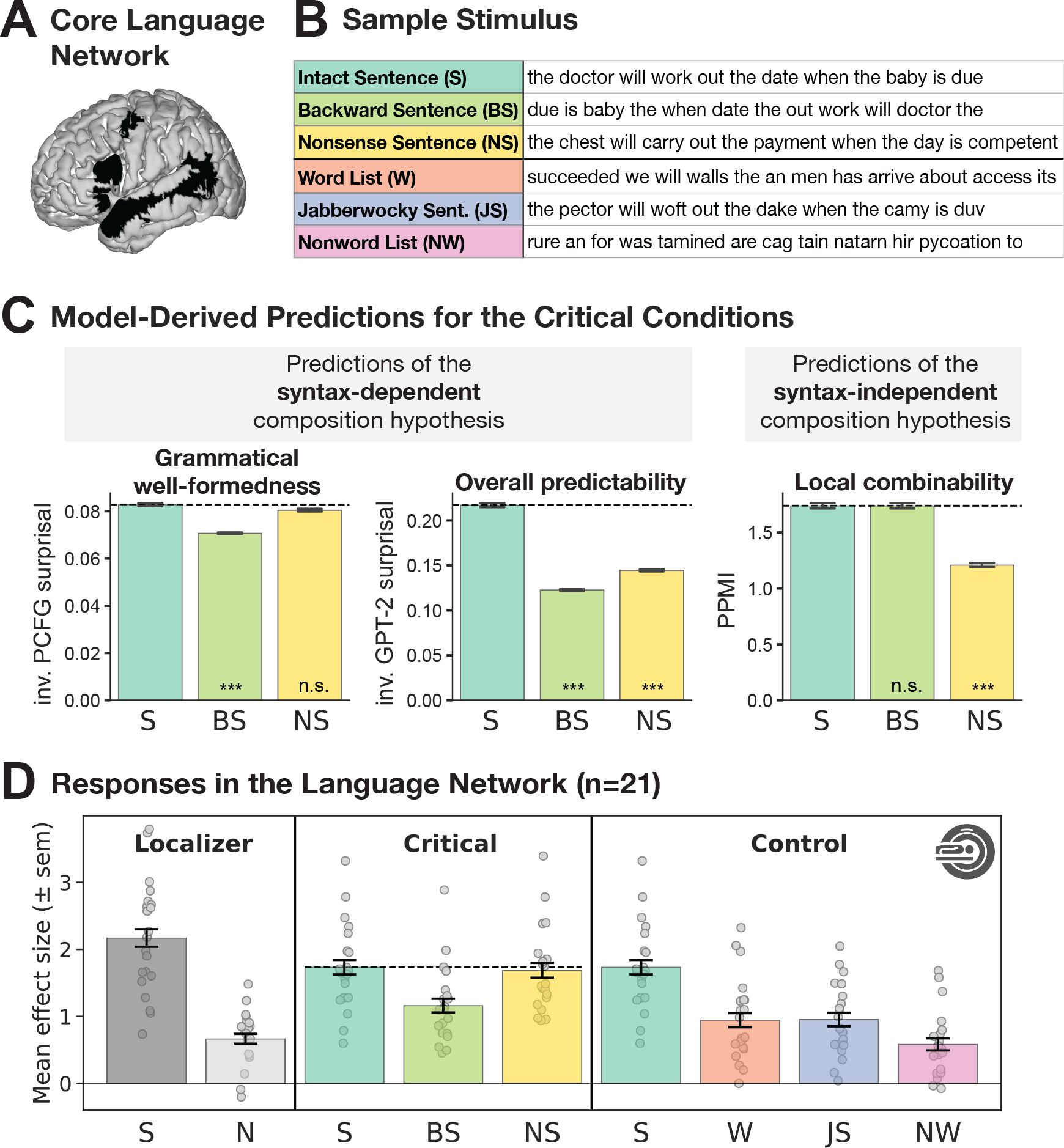
Syntax-driven meaning composition drives the language network’s response. A) The parcels that were used to functionally define the language-responsive areas in individual participants (Fedorenko et al., 2010). In each participant, the top 10% of most localizer-responsive (S>N) voxels within each parcel were taken as that participant’s region of interest. **B)** A sample item from the critical experiment. **C)** Quantitatively derived predictions for syntax-dependent vs. syntax-independent semantic composition hypotheses. The syntax-dependent panel is split up into predictions derived via structure-mediated vs. expectation-mediated incremental processing models (see <u>Discussion</u>). To match the expected direction of the neural responses in the language network, we show inverse surprisal (i.e., the reciprocal of surprisal) for the PCFG and GPT-2 models. Significant difference to the *Sentence* condition was established via post hoc pairwise t- tests, with p-values corrected for multiple comparisons using the Bonferroni procedure. **D)** Neural responses (in % BOLD signal change relative to fixation) to the conditions of the language localizer and critical and control experimental conditions within the language network (averaged across the five regions; the profiles of individual fROIs are similar, as shown in Figure 3). Dots show individual subject responses; error bars show standard errors of the mean by participants. The observed response pattern is best explained by syntax-*dependent* semantic composition (see panel C).

Next, we examined a model-derived measure to determine the degree to which a string supports *syntax-independent semantic composition*, i.e., obeys the local semantic dependency structures of naturally occurring language inputs. In other words, this measure quantifies the degree to which a processing algorithm that is syntax-independent might attempt to build complex meanings out of words within a local context based on prior linguistic experience and world knowledge. Following Mollica, Siegelman et al. (2020), we use (positive) Pointwise Mutual Information (**PMI**), an information-theoretic indicator of semantic association (Church & Hanks, 1990), and focus on positive PMI values, because negative values suggest there is no semantic dependency worth building. To derive PPMI scores, we used the procedure described in Mollica, Siegelman et al. (2020) (see **Figure SI 4** for a measure of directional PMI). In particular, for each string, we used a sliding four-word window to extract local word pairs (this is equivalent to collecting the bigrams, 1-skip-grams, and 2-skip-grams from each string). For each word pair, we then calculated its PPMI score. Probabilities were estimated using the Google *n*-gram corpus (Michel et al., 2011) and zs Python library (Smith, 2014) with Laplace smoothing (*α* = 0.1). We obtain a PPMI estimate for each word pair occurring within a four-word sliding window (see Equation (1)), and then average these estimates across all words pairs in the sentence to derive a single score per sentence.

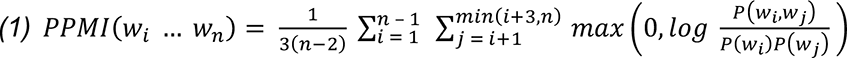

Finally, we evaluated our stimuli in a **behavioral** rating study (see **Figure 4A**). In particular, we wanted to ensure that our *Nonsense* sentences are judged as grammatically well-formed but lacking conventional meaning by naïve participants. We recruited 120 participants through Amazon’s Mechanical Turk and asked them to rate our *Nonsense* sentence stimuli, along with *Sentence* and *Word list* stimuli for comparison (192 stimuli per condition), for two features on a 1-5 Likert scale: grammatical well-formedness (1=completely ungrammatical to 5=perfectly grammatical) and semantic acceptability (1=doesn’t make any sense to 5=makes perfect sense). As part of the instructions, several examples for each concept were provided. The 576 stimuli were distributed across 4 experimental lists (144 stimuli each, 48 per condition). Each participant completed only one list. In order to counteract poor data quality on MTurk, owing to the use of bots or fake IP addresses (e.g., Chmielewski & Kucker, 2020), we restricted our task to participants with IP addresses in the United States and included only those participants in the analyses who satisfied all of the following criteria: (i) they used the full Likert scale and (ii) they did not, on average, rate *Sentence* stimuli lower than 3 in terms of grammaticality and they did not, on average, rate *Word lists* higher than 3 in terms of grammaticality. Data from 57 participants were included in the final analysis.

#### 2.2.2 Critical task procedure

For our critical fMRI experiment, each experimental list (48 stimuli x 8 conditions = 384 stimuli total) was divided into 8 subsets of 48 stimuli each (6 stimuli per condition), corresponding to 8 scanning runs. A blocked design was used, with each block consisting of 3 trials of the same condition. The trial structure was similar to that used in Mollica, Siegelman et al. (2020): the stimulus (a sequence of 12 words/nonwords) was presented one word/nonword at a time in the center of the screen for 350 ms each, in black capital letters on a white background with no punctuation. The stimulus sequence was followed by a blank screen for 300 ms, then by a memory probe word/nonword presented in blue font for 1,000 ms, and finally, by a blank screen for 500 ms, for a total trial duration of 6 s (thus, experimental blocks were 18 s in duration). When a memory probe appeared, participants were asked to determine whether the probe was the same as the last word/nonword in the sequence they just read, and to indicate their choice via pressing one of two buttons. On half of the trials, the memory probe was the same as the last word/nonword in the sequence; on the remaining trials, probes were randomly sampled correct probes from other stimuli of the same condition from a different block. The memory probe task was designed to be easy and was included to help participants stay alert. Each scanning run (consisting of 16 experimental blocks—2 blocks per condition—and 5 fixation blocks) lasted 16 ∗ 18 s + 5 ∗ 12 s = 348 s (5 min 48s).

#### 2.2.3 Localizers

##### Language network localizer task

The regions of the language network were localized using a task described in detail in Fedorenko et al. (2010) and subsequent studies from the Fedorenko lab (the task is available for download from https://evlab.mit.edu/funcloc/). Briefly, participants silently read sentences and lists of unconnected, pronounceable nonwords (each 12 word-/nonwords-long) in a blocked design. The sentences > nonwords contrast targets brain regions that that support high-level language comprehension. This contrast generalizes across tasks (e.g., Fedorenko et al., 2010; Scott et al., 2017; Ivanova et al., 2020) and presentation modalities (reading vs. listening; e.g. (Fedorenko et al., 2010; Scott et al., 2017; Chen et al., 2021; Malik-Moraleda, Ayyash et al., 2022). All the regions identified by this contrast show sensitivity to lexico-semantic processing (e.g., stronger responses to real words than nonwords) and combinatorial semantic and syntactic processing (e.g. stronger responses to sentences and Jabberwocky sentences than to unstructured word lists and nonword lists) (e.g., Fedorenko et al., 2010, 2016, 2020, 2012; I. Blank et al., 2016; Shain, Kean et al., 2024). More recent work further shows that these regions are also sensitive to sub-lexical regularities (Regev et al., 2024), in line with the idea that this system stores our linguistic knowledge, which encompasses regularities across representational grains, from phonological and morphological schemas to words and constructions (see Fedorenko et al., 2024 for a review). Stimuli were presented one word/nonword at a time at the rate of 450 ms per word/nonword. Participants read the materials passively and performed a simple button-press task at the end of each trial, which was included in order to help participants remain alert. Each participant completed 2 ∼6 min runs.

##### Multiple Demand network localizer task (relevant for some control analyses)

The regions of the Multiple Demand (MD) network (Duncan, 2010; Duncan et al., 2020) were localized using a spatial working memory task contrasting a harder condition with an easier condition (e.g., Fedorenko et al., 2011, 2013; Blank et al., 2014). The hard > easy contrast targets brain regions engaged in cognitively demanding tasks. Fedorenko et al. (2013) have established that the regions activated by this task are also activated by a wide range of other demanding tasks (see also Duncan & Owen, 2000; Hugdahl et al., 2015; Shashidhara et al., 2019; Assem et al., 2020). On each trial (8 s), participants saw a fixation cross for 500 ms, followed by a 3 x 4 grid within which randomly generated locations were sequentially flashed (1 s per flash) 2 at a time for a total of 8 locations (hard condition) or 1 at a time for a total of 4 locations (easy condition). Then, participants indicated their memory for these locations in a 2-alternative, forced-choice paradigm via a button press (the choices were presented for 1,000 ms, and participants had up to 3 s to respond). Feedback, in the form of a green checkmark (correct responses) or a red cross (incorrect responses), was provided for 250 ms, with fixation presented for the remainder of the trial. Hard and easy conditions were presented in a standard blocked design (4 trials in a 32 s block, 6 blocks per condition per run) with a counterbalanced order across runs. Each run included 4 blocks of fixation (16 s each) and lasted a total of 448 s. Each participant completed 2 runs.

#### 2.2.4 Participants

Twenty-two individuals (12 female, 10 male, mean age = 26.3 years, SD = 7.97) were recruited from MIT and the surrounding Cambridge/Boston, MA, community and paid for their participation. All were native speakers of English, had normal hearing and normal or corrected vision, and had no history of language impairment. 20 participants were right-handed, and the remaining 2 were left-handed, as determined by the Edinburgh handedness inventory (Oldfield, 1971), or self-report. All but three participants showed typical left-lateralized activation for the language localizer task (paradigm details above). Lateralization was calculated based on the number of significant (at the fixed whole-brain uncorrected voxel-level threshold of p<0.001) language-responsive voxels in the left vs. right hemispheres (LH vs. RH), using the following formula: (*LH* − *RH*)/(*LH* + *RH*), following Jouravlev et al. (2020). For the three participants, who showed right-lateralized language responses (lateralization values of -0.67, -0.64, and -0.4; individuals with values of -0.25 or below are considered right-lateralized; Jouravlev et al., 2020), we used their right-hemisphere language regions for the analyses. (We show in **Figure SI 5** that the results remain unchanged when using the left hemisphere fROIs for all participants.) One participant showed low behavioral performance on the memory probe task (*<*60% accuracy) and was excluded from the analyses, leaving a total of 21 participants. One additional participant showed low behavioral performance for the first 3 runs of the critical task (consistent with self-reported sleepiness); we excluded those runs from the analyses. All participants gave written informed consent in accordance with the requirements of MIT’s Committee on the Use of Humans as Experimental Subjects (COUHES).

#### 2.2.5 fMRI data acquisition, preprocessing, first-level modeling, and fROI definition

##### Data acquisition

Structural and functional data were collected on a whole-body 3 Tesla Siemens Prisma scanner with a 32-channel head coil at the Athinoula A. Martinos Imaging Center at the McGovern Institute for Brain Research at MIT. T1-weighted, Magnetization Prepared Rapid Gradient Echo (MP- RAGE) structural images were collected in 208 sagittal slices with 0.85 mm isotropic voxels (TR = 1,800 ms, TE = 2.37 ms, TI = 900 ms, flip = 8 degrees). Functional, blood oxygenation level-dependent (BOLD) data were acquired using an SMS EPI sequence with a 90° flip angle and using a slice acceleration factor of 3, with the following acquisition parameters: seventy-two 2 mm thick near-axial slices acquired in the interleaved order (with 10% distance factor), 2 mm × 2 mm in-plane resolution, FoV in the phase encoding (F >> H) direction 208 mm and matrix size 104 × 104, TR = 2,000 ms, TE = 30 ms, and partial Fourier of 7/8. The first 10 s of each run were excluded to allow for steady state magnetization.

##### Data preprocessing

fMRI data were analyzed using SPM12 (release 7487), CONN EvLab module (release 19b), and other custom MATLAB scripts. Each participant’s functional and structural data were converted from DICOM to NIFTI format. All functional scans were co-registered and resampled using B-spline interpolation to the first scan of the first session (Friston et al., 1995). Potential outlier scans were identified from the resulting subject-motion estimates as well as from BOLD signal indicators using default thresholds in CONN preprocessing pipeline (5 standard deviations above the mean in global BOLD signal change, or framewise displacement values above 0.9 mm; (Nieto-Castañón, 2020). Functional and structural data were independently normalized into a common space (the Montreal Neurological Institute [MNI] template; IXI549Space) using SPM12 unified segmentation and normalization procedure (Ashburner & Friston, 2005) with a reference functional image computed as the mean functional data after realignment across all timepoints omitting outlier scans. The output data were resampled to a common bounding box between MNI-space coordinates (-90, -126, -72) and (90, 90, 108), using 2mm isotropic voxels and 4th order spline interpolation for the functional data, and 1mm isotropic voxels and trilinear interpolation for the structural data. Last, the functional data were smoothed spatially using spatial convolution with a 4 mm FWHM Gaussian kernel.

##### First-level modeling

Effects were estimated using a General Linear Model (GLM) in which each experimental condition was modeled with a boxcar function convolved with the canonical hemodynamic response function (HRF) (fixation was modeled implicitly, such that all timepoints that did not correspond to one of the conditions were assumed to correspond to a fixation period). Temporal autocorrelations in the BOLD signal timeseries were accounted for by a combination of high-pass filtering with a 128 seconds cutoff, and whitening using an AR(0.2) model (first-order autoregressive model linearized around the coefficient a=0.2) to approximate the observed covariance of the functional data in the context of Restricted Maximum Likelihood estimation (ReML). In addition to experimental condition effects, the GLM design included first-order temporal derivatives for each condition (included to model variability in the HRF delays), as well as nuisance regressors to control for the effect of slow linear drifts, subject-motion parameters, and potential outlier scans on the BOLD signal.

##### fROI definition

Following prior work, we used group-constrained, participant-specific functional localization (Fedorenko et al., 2010). Namely, individual activation maps for the target contrast (here, sentences *>* nonwords) were combined with spatial ‘masks’—corresponding to broad areas within which most participants in a large, independent sample show activation for the same contrast. The masks, which were derived in a data-driven way from this independent sample of participants and are available from the lab’s website, have been used in many prior studies (e.g., Diachek, Blank, Siegelman et al., 2020; Jouravlev et al., 2019a; Shain, Blank et al., 2020). They include five regions in each hemisphere: three in the frontal cortex (two in the inferior frontal gyrus, including its orbital portion: IFGorb, IFG; and one in the middle frontal gyrus: MFG), and two in the anterior and posterior temporal cortex (AntTemp and PostTemp). Within each mask, we selected 10% of most localizer-responsive voxels (voxels with the highest t-value for the localizer contrast) following the standard approach in prior work. This approach allows to pool data from the same functional regions across participants while allowing for inter-individual variability in the precise locations of these regions. All main analyses were performed on fMRI BOLD signals extracted from these functional ROIs. For completeness, we also defined a language fROI in the angular gyrus (see **Figure SI 6, Table SI 2**). This area is activated by the language localizer contrast (e.g., Fedorenko et al., 2010) but has been shown to dissociate from the core frontal and temporal language areas (e.g., Shain, Paunov, Chen et al., 2023).

In addition to the language fROIs, we defined a set of control fROIs using the Multiple Demand network localizer. Here, individual activation maps for the hard > easy contrast were combined with a set of twenty spatial ‘masks’ (10 regions in each hemisphere), which were derived in a data-driven way from an independent sample of participants and are available from the lab’s website. The masks cover the frontal and parietal components of the MD network (Duncan, 2010, 2013) bilaterally. Similar to the language masks, these masks have been used in many prior studies (e.g., Diachek, Blank, Siegelman et al., 2020; Jouravlev et al., 2019a; Shain, Blank et al., 2020). Within each mask, we selected 10% of most localizer-responsive voxels, and the analyses were performed on fMRI BOLD signals extracted from these fROIs.

#### 2.2.5 Estimating responses in the language network to the critical conditions and statistical analyses

After defining the language fROIs (and, for control analyses, MD fROIs), we extracted the fROIs’ responses to the critical task conditions of interest. To obtain these values, we averaged BOLD responses across voxels within each language fROI in each participant to obtain a value for each condition. Because the fROIs comprising the language network are strongly functionally interconnected (Blank et al., 2014; Mineroff, Blank et al., 2018; Paunov et al., 2019), we perform the key analyses at the network level, but we also show that the response profiles are similar in each individual language fROI (**Figure 3**).

**Figure 3.**
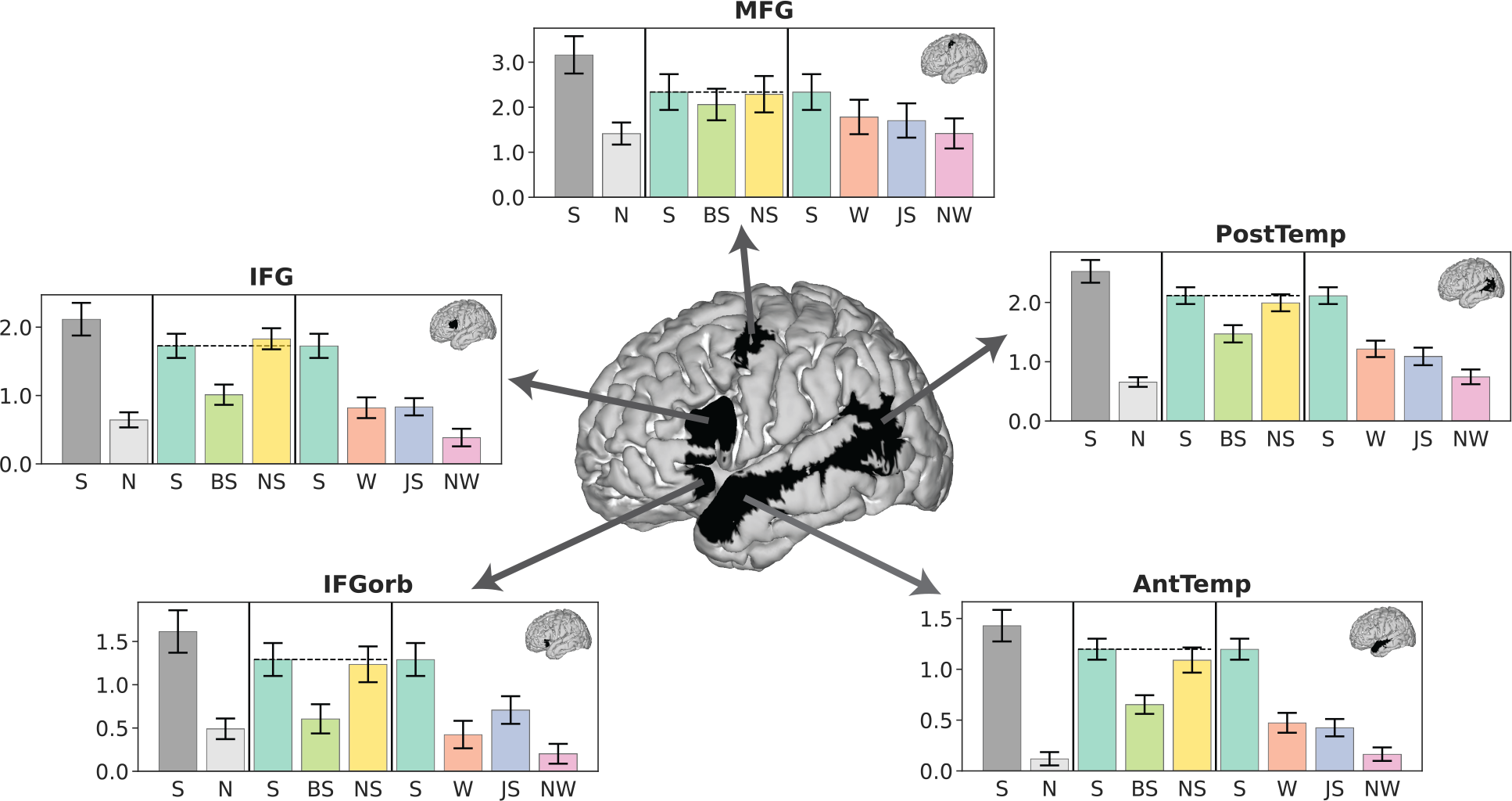
Responses in the five areas of the language network reveal the stability of the observed condition pattern. Neural responses (in % BOLD signal change relative to fixation) to the conditions of the language localizer and critical and control experimental conditions in each of the five language functional regions of interest (fROIs). IFGorb—orbital inferior frontal gyrus, IFG—inferior frontal gyrus, MFG—middle frontal gyrus, AntTemp—anterior temporal lobe, PostTemp—posterior temporal lobe. The profiles of individual fROIs are similar to each other and mirror that of the overall network response (**Figure 2D, Table SI 5**).

**Figure 4.**
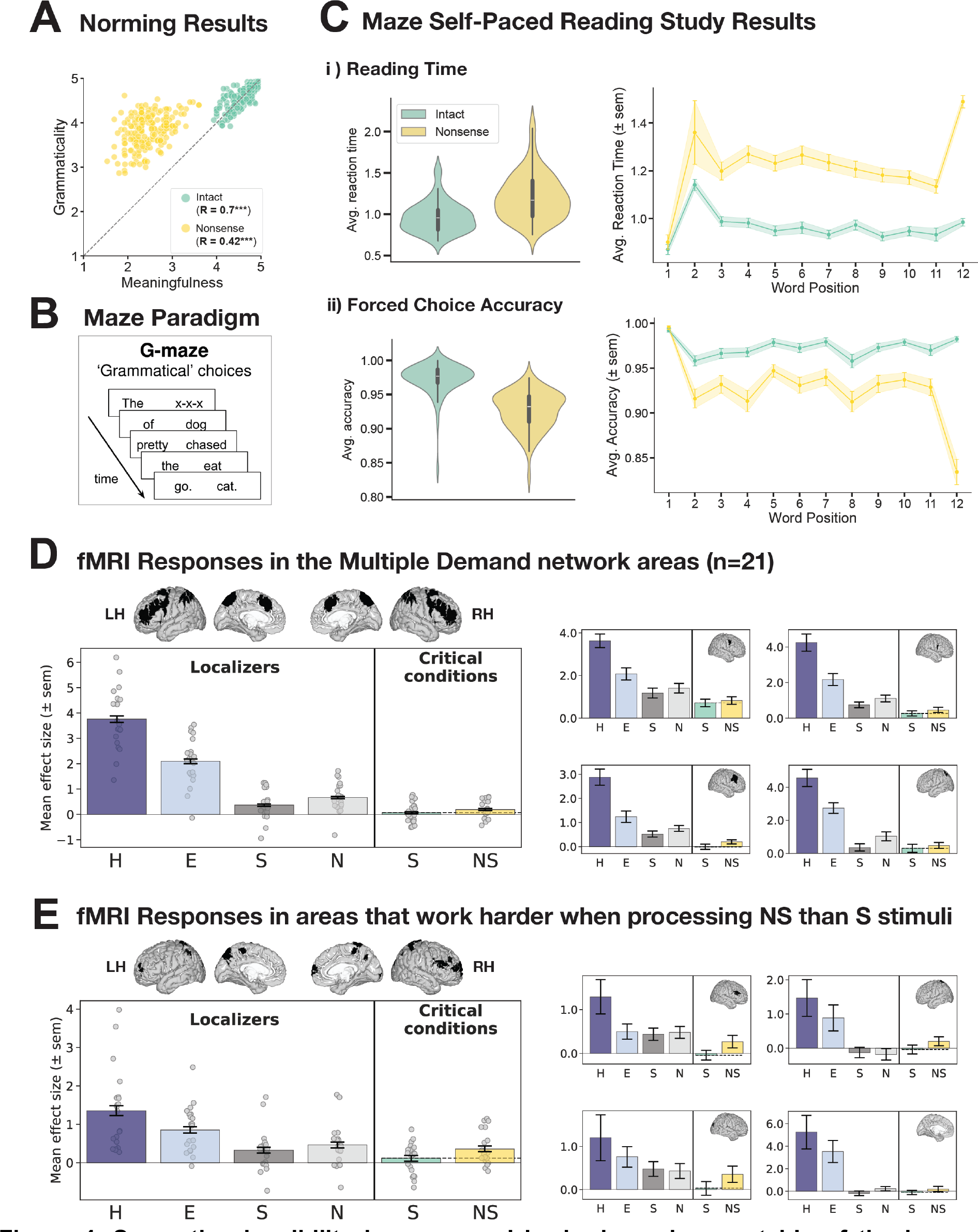
Semantic plausibility is processed by brain regions outside of the language network. Panels A-E focus on the cost associated with semantic implausibility; Panel F focuses on brain regions that respond more to plausible sentences. **A)** Behavioral norming study results for *Sentence* and *Nonsense* sentence stimuli (for details, see Section 2.2.1; for the full set of results, including *Word list* stimuli, see **Figure SI 9**). *Nonsense* sentences are rated as less meaningful, but also less grammatical than typical sentences (likely because human judgments of sentence grammaticality and meaningfulness tend to be correlated). **B)** Illustration of the Maze experimental paradigm. **C)** Maze study results for reading times (top row plots) and forced choice accuracy (bottom row plots). Violin plots show average across all positions, line plots show averages per word position. Processing *Nonsense* sentences is measurably more costly than processing typical sentences. **D)** Neural responses (in % BOLD signal change relative to fixation) to the conditions of the Multiple Demand (MD) and language localizer tasks and the *Sentence* and *Nonsense sentence* conditions of the critical experiment in the MD network. For responses to all experimental conditions and statistical analysis see **Figure SI 10**, **Table SI 6**. The black masks on the brain show the parcels that were used to define the MD areas in individual participants (see Section 2.2.3 for details). The large plot shows the average response profile across the MD network. Dots show individual subject responses; error bars show standard errors of the mean by participants. The smaller plots show the response of four sample MD ROIs; inset brains show the location of the respective parcel. The MD network as a whole shows a numerically, albeit not significantly, higher response to *Nonsense* sentences than typical sentences. **E)** Neural responses (in % BOLD signal change relative to fixation) to the conditions of the Multiple Demand (MD) and language localizer tasks and the *Sentence* and *Nonsense sentence* conditions of the critical experiment averaged across brain areas derived from a whole- brain search for areas that respond to the *Nonsense sentence* > *Sentence* contrast (n=9 regions total; responses are estimated with across-runs cross-validation) (see **Figure SI 11** for details). Dots show individual subject responses; error bars show standard errors of the mean by participants. These regions and their responses to the conditions of the MD network localizer suggest that they constitute a subset of the MD network. **F)** Neural responses (in % BOLD signal change relative to fixation) to the conditions of the Multiple Demand (MD) and language localizer tasks and the *Sentence* and *Nonsense sentence* conditions of the critical experiment averaged across brain areas derived from a whole-brain search for areas that respond to the *Sentence* > *Nonsense sentence* contrast (n=14 regions total; responses are estimated with across-runs cross- validation) (see **Figure SI 12** for details). Dots show individual subject responses; error bars show standard errors of the mean by participants. These regions and their responses to the conditions of the MD network localizer suggest that they constitute a subset of the Default Mode network.

To evaluate the statistical significance of the differences in the average change in BOLD signal across conditions, we employed a mixed-effect linear regression model with a maximal random-effect structure (Barr et al., 2013), predicting the level of response with a fixed effect and random slope for Condition, and random intercepts for Participants and ROIs (**Equation 2**).

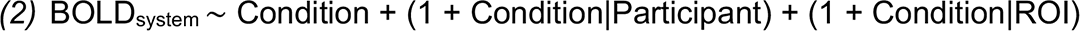

For the analyses at the ROI level, we conducted a mixed-effect linear regression model with a maximal random-effect structure, predicting the level of response with a fixed effect and random slope for Condition, and random intercepts for Participants (**Equation 3**). In both models, conditions were dummy-coded with the *Sentence* condition as the reference level.

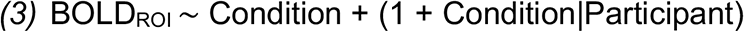

## 3. Results

We evaluate the two hypotheses laid out in the <u>Introduction</u>—the syntax-dependent semantic composition hypothesis and the syntax-independent semantic composition hypothesis—across two key analyses. First, we elicit human intuitions about incremental parsing of syntactically ill-formed linguistic inputs using a novel behavioral paradigm. We focus on linguistic inputs that have been previously argued to be processed without a detailed syntactic analysis, based on local semantic cues, and test whether such inputs could instead be reconstructed as they are processed incrementally (Section 3.1). And second, using fMRI, we examine responses in the language brain areas to stimuli designed to adjudicate between the two hypotheses (Section 3.2). In the last section, we briefly examine brain areas that process semantic plausibility, given that we find that this information is not processed within the language areas (Section 3.3).

### 3.1 Word-order scrambled stimuli that elicit a sentence-level response in the language network are amenable to real-time syntactic reconstruction

To investigate whether the invariance of the language network’s response to local word order scrambling (Mollica, Siegelman et al., 2020) may reflect the ability to parse the input following its syntactic reconstruction, rather than shallow, semantics-based comprehension, we probed people’s ability to reconstruct syntactically ill-formed inputs using SynReco—a novel behavioral paradigm (**Figure 1A**; for a validation of the paradigm, see **Figure SI 7**). We used the conditions from the Mollica, Siegelman et al.’s study, including intact sentences (*Intact*), sentences with local word order scrambling (with 1, 3, 5, or 7 local swaps; *Scrambled{1,3,5,7}*), and sentences where the word order is scrambled in a way that destroys local dependencies (*Scrambled_LowPMI*) (**Figure 1B**). We also added a critical new condition, which retains all local dependencies of the *Intact* sentence (**Figure 1D**) but should make syntactic reconstruction in real time challenging: sentences presented in reverse order (*Backward*).

In the analyses, we focus on the final submitted word orders, i.e., the orders submitted at the last time step. To measure a string’s grammatical well-formedness, we leverage PCFG parser surprisal. PCFG surprisal estimates do not encode surface-level patterns of word co-occurrence directly and instead rely on the structured syntactic representations of stimuli (see also Shain, Blank et al., 2020). To establish a baseline, we first quantified the grammaticality of our set of experimental stimuli across conditions (**Figure 1C, hatched bars**). We find that, as expected, PCFG surprisal increases numerically with every increase in word-order degradation and is highest for the *Scrambled_LowPMI* and *Backward* conditions (see **Table SI 3, Experimental Items,** for pairwise Tukey’s HSD (honestly significant difference) significance testing).

Next, we quantified the grammaticality of participants’ reconstructions of these stimuli across conditions (**Figure 1C, solid bars**). We find that for all conditions where word order was perturbed, participants managed to make the stimuli more grammatically well-formed (**Figure 1C, D**), even though every additional word swap was associated with a significant decrease in the ability to reconstruct the original sentence verbatim (see **Figure 1E**, **Table SI 4**), consistent with the offline reconstruction results reported in Mollica, Siegelman et al. (2020). Critically, reconstructions of stimuli with local word order swaps, for which Mollica, Siegelman et al. report sentence-level responses in the language network (*Scrambled{1,3,5,7}*), mostly do not significantly differ from one another (see **Table SI 3** for all pairwise comparisons) but are all significantly more well-formed than reconstructions of the *Scrambled_LowPMI* stimuli where local dependencies are destroyed and for which responses in the language network are low (all ps < 0.001, **Table SI 3**). In other words, this paradigm effectively captures behavioral patterns that plausibly underlie the observed pattern of responses in the language areas, as reported by Mollica, Siegelman et al. (2020). The stimuli in the *Backward* condition pattern with the stimuli in the *Scrambled_LowPMI* condition (**Table SI 3**). If the ability to parse the input (following incremental reconstruction, if need be, for syntactically ill-formed inputs) is indeed tied to the level of response to those inputs in the language brain areas, we would expect the *Backward* condition to pattern with the *Scrambled_LowPMI* condition (i.e., to elicit a relatively low response). This is indeed what we find, as discussed next.

### 3.2 Syntax-driven, not syntax-independent, semantic composition drives the language network’s response

In Section 3.1, we established that stimuli with local word order scrambling that have previously been shown to elicit a sentence-level response in the language network are amenable to real-time syntactic reconstruction, whereas stimuli with more severe word order scrambling are not. One of the latter conditions, the *Scrambled_LowPMI* condition, where local dependencies are destroyed, was shown by Mollica, Siegelman et al. (2020) to elicit a relatively low response in the language network. The *Backward* condition is similar to the *Scrambled_LowPMI* condition in the difficulty of online syntactic reconstruction but importantly, local dependencies are preserved. This condition therefore constitutes a critical test for the syntax-independent semantic composition hypothesis: if the language network’s response is driven by shallow/associative processing operations, which are guided by local, word-order-independent semantic relationships, then this condition should elicit a strong response, similar to these brain regions’ response to well-formed sentences (see **Figure 2C** for quantitative evidence of similar local combinability; see Section 2.2.1 for details of this measure). If, on the other hand, language comprehension is syntax-driven, then this condition should elicit a low response in the language network given that syntactic structure cannot be reconstructed incrementally and thus the input cannot be parsed (see also **Figure 2C** for quantitative evidence of lower syntactic well-formedness based on PCFG surprisal; see Section 2.2.1 for details of this measure).

Another condition that helps distinguish between our two hypotheses are grammatically well-formed sentences where building complex meanings is impeded by semantic implausibility (*Nonsense* sentences). If language comprehension is syntax-driven, this condition should elicit a strong response in the language network (see **Figure 2C**, leftmost panel, for quantitative evidence of syntactic well-formedness based on PCFG surprisal); in contrast, if language comprehension is shallow/associative, then this condition should elicit a low response because the lack of plausible semantic dependencies should discourage complex meaning construction (see **Figure 2C**, rightmost, for quantitative evidence of local combinability based on PPMI).

In fMRI, we collected responses to *Backward* sentences and *Nonsense* sentences, as well as to well-formed sentences and three linguistic control conditions that are commonly used to probe computations related to lexical access vs. structure building and that can be helpful for interpreting the responses to the critical conditions—*Word Lists*, *Jabberwocky Sentences,* and *Nonword Lists* (e.g., Fedorenko et al., 2010, 2012, 2016; Shain, Kean et al., 2024).

Condition-level effects in the language network are reported in **Figure 2D**. First, replicating much prior work (e.g., Fedorenko et al., 2010, 2012, 2016; Shain, Kean et al., 2024), we found a pattern whereby the *Intact Plausible Sentence (Sentence)* condition elicited the strongest response in the language areas, *Word Lists* and *Jabberwocky Sentences* elicited a lower response, and *Nonword Lists* elicited the lowest response (**Figure 2D, control conditions**). Second and critically, we found that the *Backward Sentence* condition elicited a significantly lower response relative to the *Sentence* condition, whereas the *Nonsense Sentence* condition elicited a response that was similar in magnitude to and statistically indistinguishable from that observed for *Sentences* (**Table 1**, **Figure 2D, critical conditions**; see **Figure SI 5** for evidence that the results hold when using left-hemisphere language fROIs for all participants, including those with right-lateralized language activations). The results also held—both qualitatively and statistically—for each language ROI separately (**Figure 3** and **Table SI 5**), and in individual participants (**Figure SI 8**), evidencing their robustness. (Even if the small numerical difference in magnitude between the language network’s response to *Sentence* vs. *Nonsense Sentence* conditions—driven by the two temporal fROIs (see **Figure 3**)—becomes statistically significant in a larger dataset, this difference is too small to be practically significant (Sullivan & Feinn, 2012): the *Nonsense Sentence* condition’s magnitude is ∼97% of the *Sentence* condition’s magnitude; cf. the *Backward Sentence* condition, which is only ∼67% of the *Sentence* condition’s magnitude.)

**Table 1:** Results of mixed-effects linear regression for fMRI responses within the language network. Stimulus type was dummy-coded with *Intact sentence* as the reference level. **\***Denotes significant difference.

|  | Estimate | Est. error | 95% CI |  |
| --- | --- | --- | --- | --- |
| Intact sentence | 1.45* | 0.53 | 0.43 | 2.54 |
| Backward sentence vs. Intact sentence | -0.55* | 0.13 | -0.80 | -0.30 |
| Nonsense sentence vs. Intact sentence | -0.08 | 0.15 | -0.38 | 0.22 |
| Word list vs. Intact sentence | -0.77* | 0.13 | -1.02 | -0.52 |
| Jabberwocky sentence vs. Intact sentence | -0.75* | 0.13 | -1.00 | -0.50 |
| Nonword list vs. Intact sentence | -1.09* | 0.14 | -1.36 | -0.82 |

The observed pattern of neural responses challenges the claim from Mollica, Siegelman et al. (2020) that the ability to form dependencies among nearby words is necessary and sufficient to elicit a sentence-level BOLD response in the language network. In particular, the network’s strong response to *Nonsense* sentences suggests that local combinability is *not necessary* to drive the network (or, at least, that the critical aspects of local combinability have to do with *syntax* rather than meaning); and the network’s low response to *Backward* sentences suggests that local combinability is *not sufficient* to elicit a sentence-level response. Instead, the findings support the hypothesis whereby the language network supports syntax-dependent semantic composition.

### 3.3 Semantic plausibility is evaluated outside of the language network

In Section 3.2, we established that *Nonsense* sentences elicit a BOLD response in the language network that is similar in magnitude to that elicited by meaningful sentences, which suggests that this particular brain system is relatively insensitive to semantic plausibility information (**Figure 2D**). However, behaviorally, *Nonsense* sentences are rated as less meaningful and less grammatical than typical plausible sentences (**Figure 4A**), so there must be a cost to processing them (see Marslen-Wilson & Tyler, 1975, 1980 for earlier evidence). In an effort to understand these effects better, we quantified this cost using self-paced reading estimates and searched for brain regions outside of the language network that are sensitive to semantic plausibility.

We used Boyce et al.’s (2020) version of the Maze self-paced reading paradigm (Freedman and Forster, 1985; Forster et al., 2009), where participants read stimuli word-by-word by successively choosing the likely next word over a contextually inappropriate distractor word in a forced-choice design (**Figure 4B**; Boyce et al., 2020; Wilcox et al., 2021) (for experiment details see <u>Supplementary Methods</u>). We find that processing *Nonsense* sentences is associated with significantly longer reading times (paired-samples t-test, *Nonsense* vs. *Sentence*: t=14.3; p<0.001) and significantly lower choice accuracy (paired-samples t-test, *Nonsense* vs. *Sentence*: t=-18.72; p<0.001) (**Figure 4C**, violin plots). Furthermore, the processing cost is immediate and persistent, manifesting across all word positions where a distractor word is present (i.e., starting with the second word), with the highest processing cost observed at the final position, i.e., at the word that co-occurred with a sentence-final period (**Figure 4C**, line plots).

What brain system might process the cost associated with semantic implausibility during language comprehension? First, we examined the responses to our critical conditions within the domain-general Multiple Demand (MD) network, which is recruited whenever humans solve demanding cognitive tasks (Duncan, 2010, 2013; Duncan et al., 2020). Although this network does not contribute to core linguistic computations related to lexical access and syntactic structure building for naturalistic linguistic inputs, in the absence of external task demands (Diachek, Blank, Siegelman et al., 2020; Shain, Blank et al., 2020; see Fedorenko & Shain, 2021 for review), it has been implicated in the processing of some perceptually and linguistically degraded linguistic inputs (e.g., Fedorenko, 2014; Kuperberg et al., 2003). For example, much prior work has reported stronger responses in this network to word lists and nonword lists than to sentences (Mineroff, Blank et al., 2018; Diachek, Blank, Siegelman et al., 2020), and Mollica, Siegelman et al. (2020) found that some regions within this network exhibit an increase in activity for degraded-word-order stimuli. We replicate these effects—for both the language localizer conditions and the conditions of the critical experiment—and extend them to a new condition (sentences presented backwards) (**Figure SI8**). For our critical contrast, between *Nonsense* and typical, plausible *Sentences*, the MD network as a whole showed a small numerically, albeit not significantly, higher response to *Nonsense* sentences (**Figure 4D**, large plot; **Table SI 6**); a few regions, mostly in the right hemisphere, showed this effect most clearly, although the magnitude of the effect is small even in those regions (**Figure 4D**, small plots).

To test whether areas elsewhere in the brain may show a higher response to *Nonsense* sentences, we additionally performed a group-constrained subject-specific (GSS) whole-brain analysis (Fedorenko et al., 2010; Julian et al., 2012). This analysis is similar to the standard random-effects group analysis in fMRI but allows for inter-individual variability in the precise locations of functional areas, which is known to exist in the association cortex (Frost & Goebel, 2012; Tahmasebi et al., 2012), yielding higher sensitivity (Nieto-Castañón & Fedorenko, 2012). We found a few areas where most participants showed a small *Nonsense sentences > Plausible sentences* effect (quantified with an across-runs cross-validation procedure); however, the topography of these regions and their responses to the MD network localizer conditions suggest that they constitute a subset of the MD network (**Figure 4E**; **Figure SI 11**). Thus, the cognitive cost associated with the processing of semantically implausible sentences is carried, in a distributed fashion, by parts of the domain-general MD network.

In addition to brain regions that are sensitive to the cost of semantic implausibility, there must also exist brain regions that respond more when the sentences convey plausible meanings that can be related to our general world knowledge. Therefore, we also performed a GSS whole-brain analysis for the *Sentences* > *Nonsense sentences* contrast. This search yielded a few regions that showed reliably greater responses to this contrast (quantified with across-runs cross-validation). Some of these regions lie in close spatial proximity to the language network, but their profiles are clearly functionally distinct (**Figure 4F; Figure SI 12**; see Ivanova, 2022 for related evidence). In particular, these regions resemble the profile of the Default Mode network (Buckner & DiNicola, 2019): they deactivate to the demanding spatial working memory task (and more so for the more demanding condition) and they respond weakly or not at all to the language localizer contrast (e.g., Mineroff, Blank et al., 2018; Braga et al., 2020; DiNicola et al., 2023). These findings align with claims that the Default Mode network supports aspects of semantic processing (Binder et al., 1997; Wirth et al., 2011; Jackson et al., 2016; Baldassano et al., 2018).

## 4. Discussion

Language inputs typically adhere to grammatical rules and express propositions that align with our knowledge of the world. But to what degree does comprehension rely on syntactic vs. semantic cues? We pushed the grammaticality and meaningfulness of linguistic inputs to their logical extremes—obliterating parsability or meaningfulness—and found support for strong reliance on the syntactic mode of processing, rather than on shallow/semantics-based processing, in the language network. Below, we discuss these findings further and contextualize them with respect to past work.

### 4.1 The language network’s core computation is syntax-dependent semantic composition

We found that the language network is engaged *whenever parsing is possible,* either on the stimuli in their original form, or following reconstruction. Mollica, Siegelman et al. (2020) reported a relative insensitivity of the language network’s (Fedorenko et al., 2024) response to local word-order scrambling. They attempted to rule out a reconstruction interpretation, but their reconstruction task may not provide a good measure of *real-time interpretability*. Indeed, using an incremental reconstruction task, we found that the locally-scrambled sentences are amenable to real-time reconstruction. Thus, the relative insensitivity of the language network to local scrambling plausibly reflects the parsability of these stimuli following incremental reconstruction of the sentence structure, not the fact that they are processed in a shallow way, based on semantic cues.

We further found that sentences presented backwards pose difficulty for real-time reconstruction, and elicit a low response in the language areas. Thus, the local combinability of words—which is preserved in these stimuli—does not invariably lead to a sentence-level response in the language areas, contra Mollica, Siegelman et al. (2020). Without syntactic (e.g., word order) cues, our prior linguistic experience and world knowledge cannot effectively guide interpretation. At the same time, we found that nonsensical sentences elicit as high a response in the language areas as plausible sentences in spite of the fact that our prior linguistic experience and world knowledge should guide our parser away from attempts to build syntactic structure. We therefore conclude that the critical aspects of local combinability that the language network cares about have to do with *syntactic* combinatoriality.

### 4.2 Syntactic reconstruction is supported by the domain-general Multiple Demand network

The language areas responded weakly to backward sentences; in contrast, the areas of the domain-general Multiple Demand (MD) network (Duncan, 2010, 2013; Duncan et al., 2020) responded more strongly to backward sentences than to well-formed sentences (**SI-9**). Together with Mollica, Siegelman et al.’s (2020) findings, these results suggest that syntactic reconstruction costs are carried by the MD network, which supports diverse computations during cognitively demanding tasks (Duncan & Owen, 2000; Fedorenko et al., 2013; Hugdahl et al., 2015; Shashidhara et al., 2019; Assem et al., 2020). Thus, although the MD network does not appear to support computations related to lexical access and syntactic parsing for well-formed linguistic inputs (e.g., Diachek, Blank, Siegelman et al., 2020; see Fedorenko & Shain, 2021 for review), it evidently supports syntactic reconstruction of corrupt inputs. Syntactically corrupt language stimuli may therefore provide a fertile testbed for probing inter-network interactions using temporally-resolved methods, like intracranial recordings. For example, the MD network must show strong stimulus-related activity for stimuli that require reconstruction (cf. Blank & Fedorenko, 2017), and it must be engaged early on, or even simultaneously with the language network given the speed of language comprehension.

### 4.3 Syntax versus other information sources during language comprehension

The similarly strong response of the language network to plausible sentences and grammatical strings that do not express conventional meanings aligns with the human ability to understand novel sentences, including those that describe unusual events or scenarios that are hypothetical, counterfactual, or fictitious. This is the generative power of language: our language system must be able to apply its computations to any syntactically well-formed sequence of words. Nevertheless, language statistics also reflect the distributional properties of objects and events in the world (Mikolov et al., 2013; Pennington et al., 2014; Roads & Love, 2020; Abdou et al., 2021; Kauf, Ivanova et al., 2023), which might lead to the expectation of sensitivity of the language brain areas to the plausibility of sentence meanings. Furthermore, a large body of psycholinguistic work has demonstrated that language comprehension is affected by diverse information sources other than syntax (e.g., MacDonald et al., 1994; Tanenhaus et al., 1995), including world knowledge (e.g., Hagoort et al., 2004; McRae & Matsuki, 2009).

Debates about whether syntactic information is processed separately from other information that can affect language comprehension raged in the psycholinguistic literature in the 1980s-2000s (for summaries, see e.g., Gibson & Pearlmutter, 1998; Clifton Jr. et al., 2003). Recent advances in our understanding of the neural architecture of language allow us to revisit these questions through a new lens. In particular, given the separability of the language-selective network from other brain systems (Fedorenko et al., 2024), this question can be recast as *whether non-syntactic information gets represented and processed in the language network*. We have already learned for example, that in line with the behavioral evidence of quick integration of lexical cues during interpretation (e.g., MacDonald et al., 1994), there doesn’t appear to be spatial segregation between neural populations that process syntactic structure and those that process word meanings (e.g., Fedorenko et al., 2020; Hu, Small et al., 2023; Shain, Kean et al., 2024). In contrast, information like gestures, facial expressions, and prosody appear to be processed by brain areas that are distinct from the language network (e.g., Deen et al., 2015; Pritchett et al., 2018; Jouravlev et al., 2019b; Regev et al., in prep.). Similarly, discourse-level structure appears to be processed in distinct, non-language-selective systems (Ferstl & von Cramon, 2001; Kuperberg et al., 2006; Ferstl et al., 2008; Lerner et al., 2011; Jacoby & Fedorenko, 2020; see Fedorenko et al., 2024 for a review).

What about world knowledge/plausibility? The fact that sentence plausibility does not affect the computations of the language network suggests that distinct brain systems support linguistic decoding vs. evaluating the meanings with respect to world knowledge. Prior evidence supports a dissociation between linguistic and general-semantic processing: pre-verbal infants and individuals with aphasia (linguistic deficits) can understand the world (e.g., Hirsh-Pasek & Golinkoff, 2010; Spelke, 2023; Chertkow et al., 1997; Saygın et al., 2004; Antonucci & Reilly, 2008; Warren & Dickey, 2021) and make sophisticated judgments about objects and events (e.g., Varley & Siegal, 2000; Dickey & Warren, 2015; Colvin et al., 2019; Ivanova et al., 2021; Benn, Ivanova et al., 2023), and distinct brain areas are activated by i) linguistic event descriptions selectively versus ii) both linguistic and non-linguistic events (Baldassano et al., 2018; Wurm & Caramazza, 2019; Ivanova et al., 2021). In line with the latter, we find that sentence plausibility is processed in brain regions that are distinct from the language-selective regions, although located in proximity to them (**Figure 4F**; **Figure SI 12**; see Ivanova et al., 2022 for related evidence).

Spatial separability between language areas and semantic areas (or areas that process eye gaze and gestures) need not imply temporal staging of linguistic decoding vs. the processing of non-linguistic information. Many brain areas may work in parallel and exchange information on a fast timescale during incremental comprehension (although the details of these parallel computations and inter-areal/inter-network interactions remain poorly understood). However, it is also possible that—at least with respect to world knowledge—some temporal staging is required: after all, to understand *that* a sentence is implausible, you need to first decode its meaning.

### 4.4 Limitations, future directions, and open questions

None of the current methods in language research directly tap into the parsing operations that humans engage in. The behavioral paradigm we developed to elicit incremental intuitions about parsing is a step in the right direction, but future work should better align the timing of the task with real-life comprehension, incorporate working memory constraints (Christiansen & Chater, 2016), and expand the repertoire of reconstruction operations (Gibson et al., 2013). Future work should also test whether the current results generalize to flexible-word-order languages, and investigate comprehension of syntactically degraded stimuli in longer contexts given past evidence of contextual influences (e.g., Spivey & Tanenhaus, 1998; Chen et al., 2023).

Interpretation-wise, we discuss our results in terms of syntax-driven semantic composition, but they could alternatively be construed in terms of prediction (e.g., Kuperberg & Jaeger, 2016). In other words, perhaps the language network gets engaged whenever we process linguistic stimuli that are predictable to some degree. The strong response to *Nonsense* sentences rules out a version of this hypothesis that has to do with *overall* predictability (**Figures 2-3**; see **Figure SI 13** for additional evidence), but our results are compatible with *purely syntactic* predictability. In fact, it is unclear whether syntactic integration and syntactic predictability could even in principle be distinguished given that stimuli where syntactic integration is possible are necessarily characterized by some degree of syntactic predictability.

Future studies should further illuminate the contributions of the Multiple Demand (MD) network to syntactic reconstruction. For example, these findings could be connected to the ERP research that has interpreted the domain-general P600 component (e.g., Osterhout & Holcomb, 1992; Patel, 2003; Núñez-Peña & Honrubia-Serrano, 2004; Cohn et al., 2012) as indexing error correction (Ryskin, Stearns et al., 2021). Our findings also make predictions about the processing of syntactically corrupt inputs in children and older adults (given that the MD network is slow to develop (Fiske & Holmboe, 2019; Schettini et al., 2023) and shows clear age-related decline (Reuter-Lorenz et al., 2000; Mitchell et al., 2023; Wu & Hoffman, 2023)). If the MD network cannot effectively support syntactic reconstruction in these populations, we should observe larger effects of syntactic degradation on the language network’s responses, and greater reliance on semantic plausibility in interpreting corrupt inputs (e.g., Beese et al., 2019).

Future work should also investigate a) the time-course and nature of the language network’s interaction with non-language-specific systems during incremental comprehension; b) the criteria that determine whether a stimulus is in-domain vs. out-of-domain for the language network (e.g., a locally scrambled sentence vs. a list of unconnected words) (see Shain, Kean et al., 2024 for further discussion); and c) the nature of linguistic representations and computations. Regarding the latter, we have here talked about parsing in terms of symbolic operations (Pullum & Gazdar, 1982; Joshi, 1985; Steedman, 2000; Chomsky, 2014), but neural network language models suggest that explicit encoding of symbolic components may not be necessary: these models reliably encode sentence structure in their embeddings (e.g., Hewitt & Manning, 2019; Sinha et al., 2021; Tucker et al., 2021; Eisape et al., 2022; see Pavlick, 2023 for discussion). Whether syntactic representations in the human brain might similarly be encoded without explicit symbolic representations, and whether/how symbolic-like representations emerge in neural network architectures remain exciting questions for future work.

## Data and code availability

Data and code will be made publicly available upon publication.

## Acknowledgments

We would like to acknowledge the Athinoula A. Martinos Imaging Center at the McGovern Institute for Brain Research at MIT and its support team (Steve Shannon and Atsushi Takahashi). We thank Cory Shain, Rachel Ryskin, Anya Ivanova, Richard Futrell, and Peng Qian for helpful comments, and Maria Ryskina for help with the fMRI data collection. CK and this work was partially supported by the K. Lisa Yang Integrative Computational Neuroscience (ICoN) Center at MIT and the MIT Quest for Intelligence. EF was supported by NIH awards R01-DC016607, R01- DC016950, and U01-NS121471, as well as by research funds from the McGovern Institute for Brain Research, the Brain and Cognitive Sciences department, the Simons Center for the Social Brain, and the Middleton professorship.

## Supplementary Information

### Supplementary Figures

**Figure SI 1.**
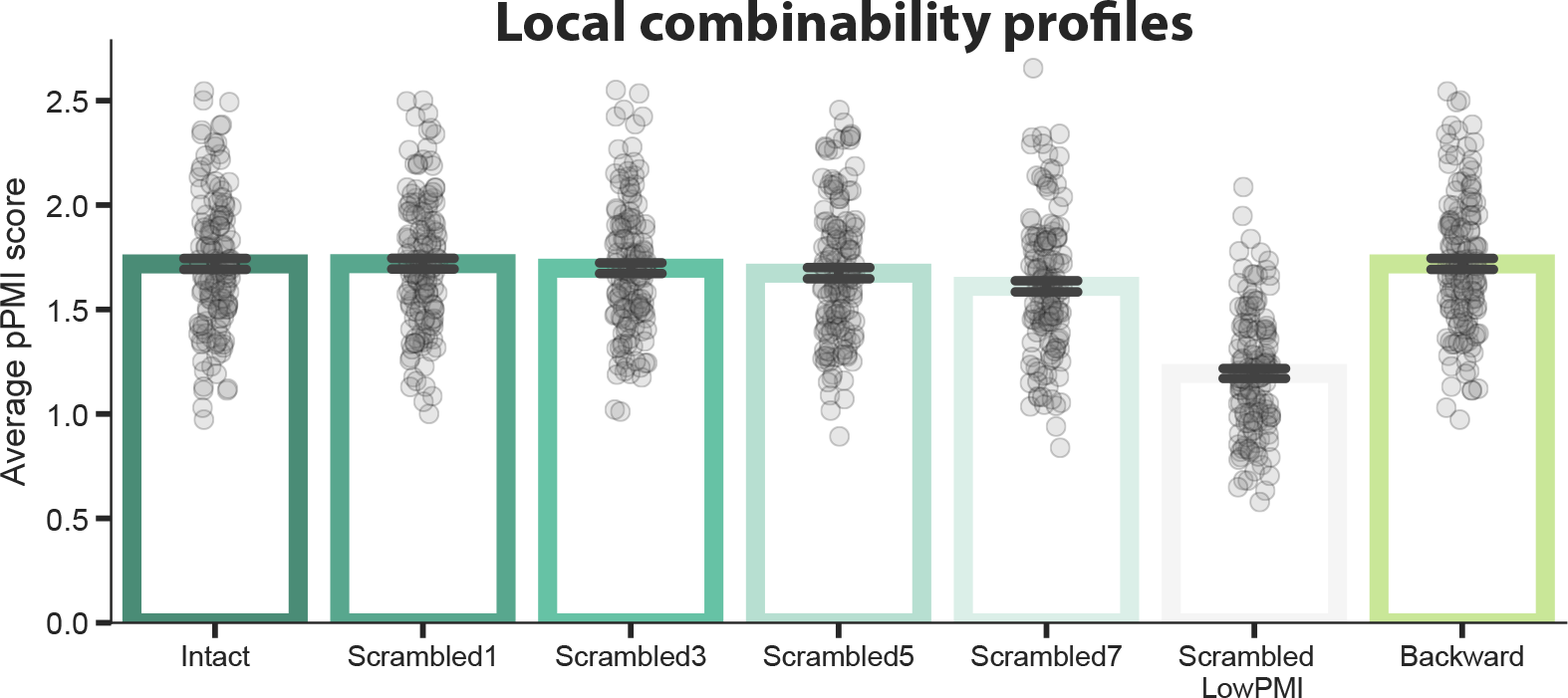
PMI calculation for the full set of materials used in Mollica, Siegelman et al.’s (2020) experiment 1, and the full set of our stimuli from the *Backward* condition.

**Figure SI 2.**
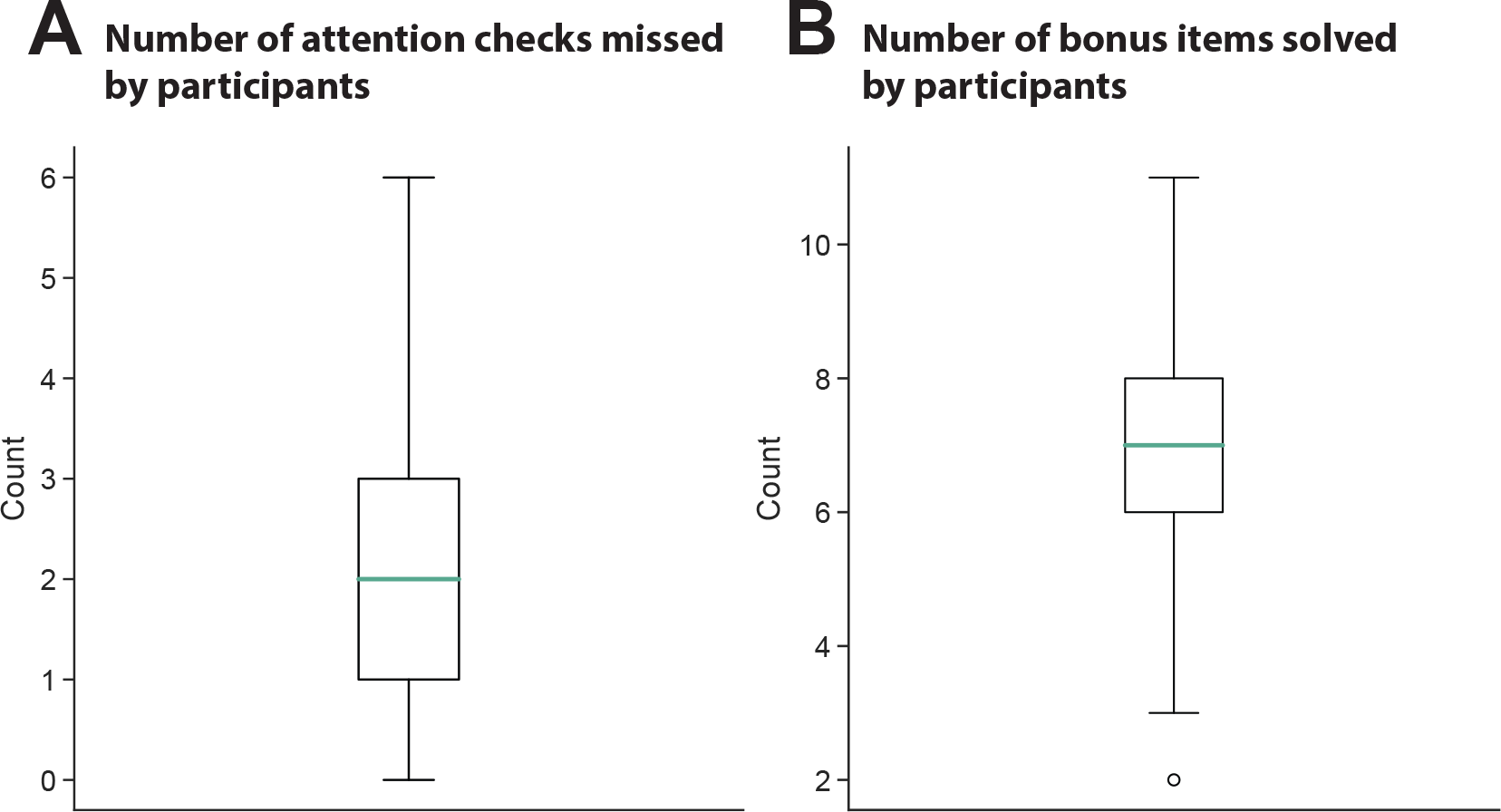
Participant exclusion criteria used for the behavioral reconstruction experiment. We used the error distributions for attention check and bonus items to determine the thresholds for exclusion. We excluded the lowest participants in the highest quartile for the number of attention checks missed, and in the lowest quartile for the number of bonus items solved.

**Figure SI 3.**
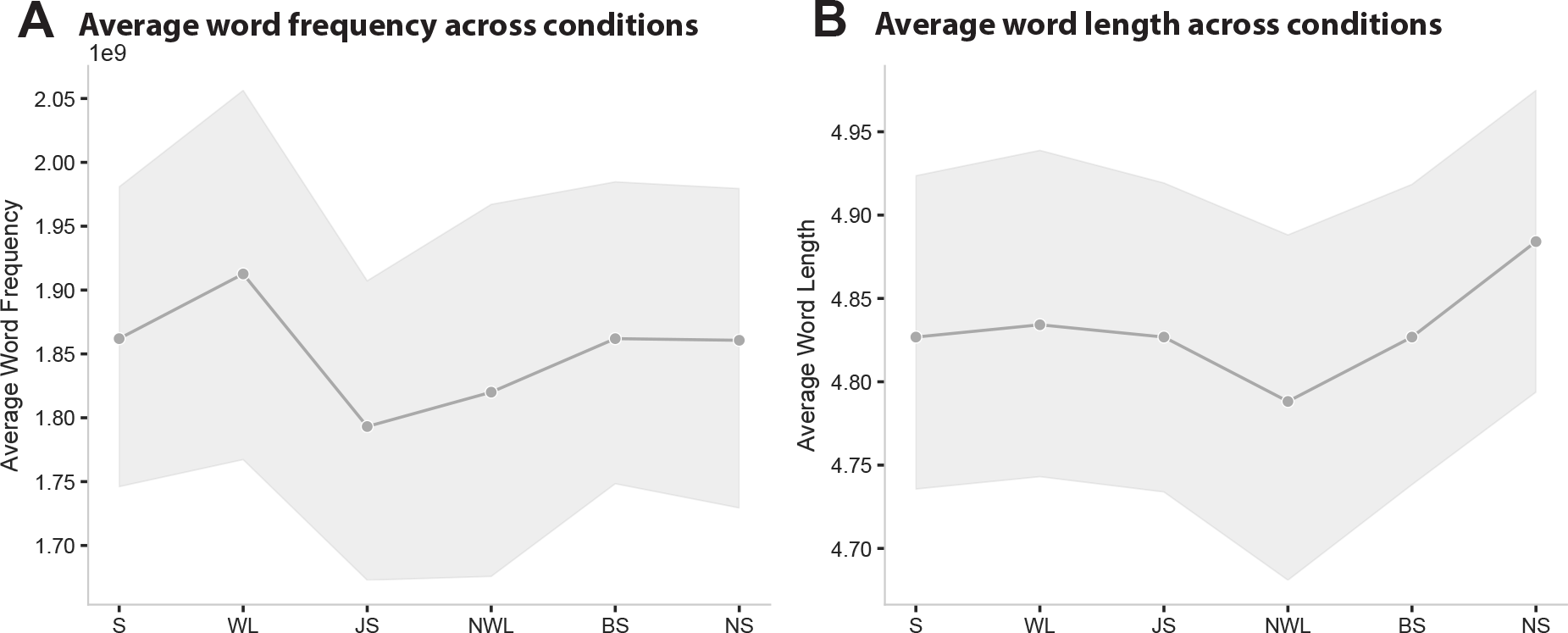
Low level controls for our experimental conditions. Stimuli are roughly matched in average word frequency and in average word length. Frequency was operationalized as the log of the number of occurrences of the word/phrase in the 2012 Google NGram corpus. Laplace smoothing was applied prior to taking the log. Word length indicates the average number of characters per word across the stimuli in a given condition.

**Figure SI 4.**
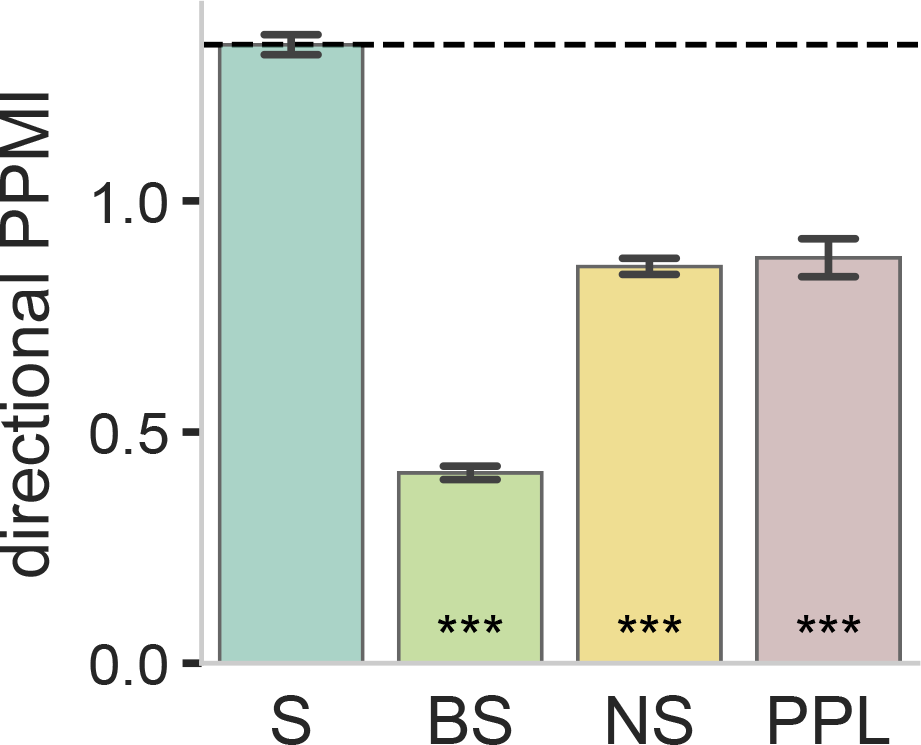
Hypothesis profile derived from directional PPMI measure. Predictions derived from a variation of the PPMI model described in <u>2.2.1 Critical task design and materials</u> that considers word order, i.e., calculates the co-occurrence of the *ordered* bigram *w*_*i*_*w*_*j*_. Significant difference to the *Sentence* condition was established via post hoc pairwise t-tests, with p-values corrected for multiple comparisons using the Bonferroni procedure. The ordered PPMI measure underpredicts the language network activity in response to *Nonsense* stimuli (and overpredicts the response to *Predictable Phrase Lists* (**Figure SI 13**)).

**Figure SI 5.**
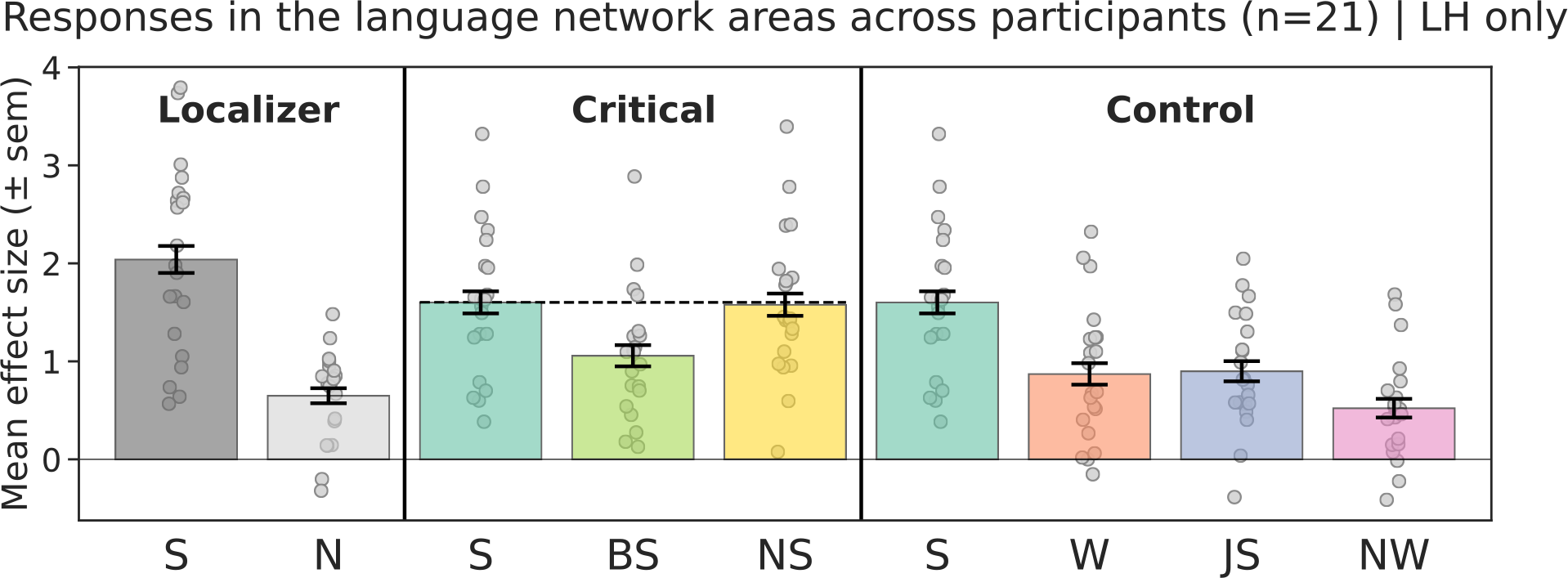
fMRI results for left-hemisphere language regions only. Neural responses (in % BOLD signal change relative to fixation) to the conditions of the language localizer and critical and control experimental conditions within the language network (averaged across all five fROIs) when including the LH (instead of the RH) language fROIs for the right-lateralized participants.

**Figure SI 6.**
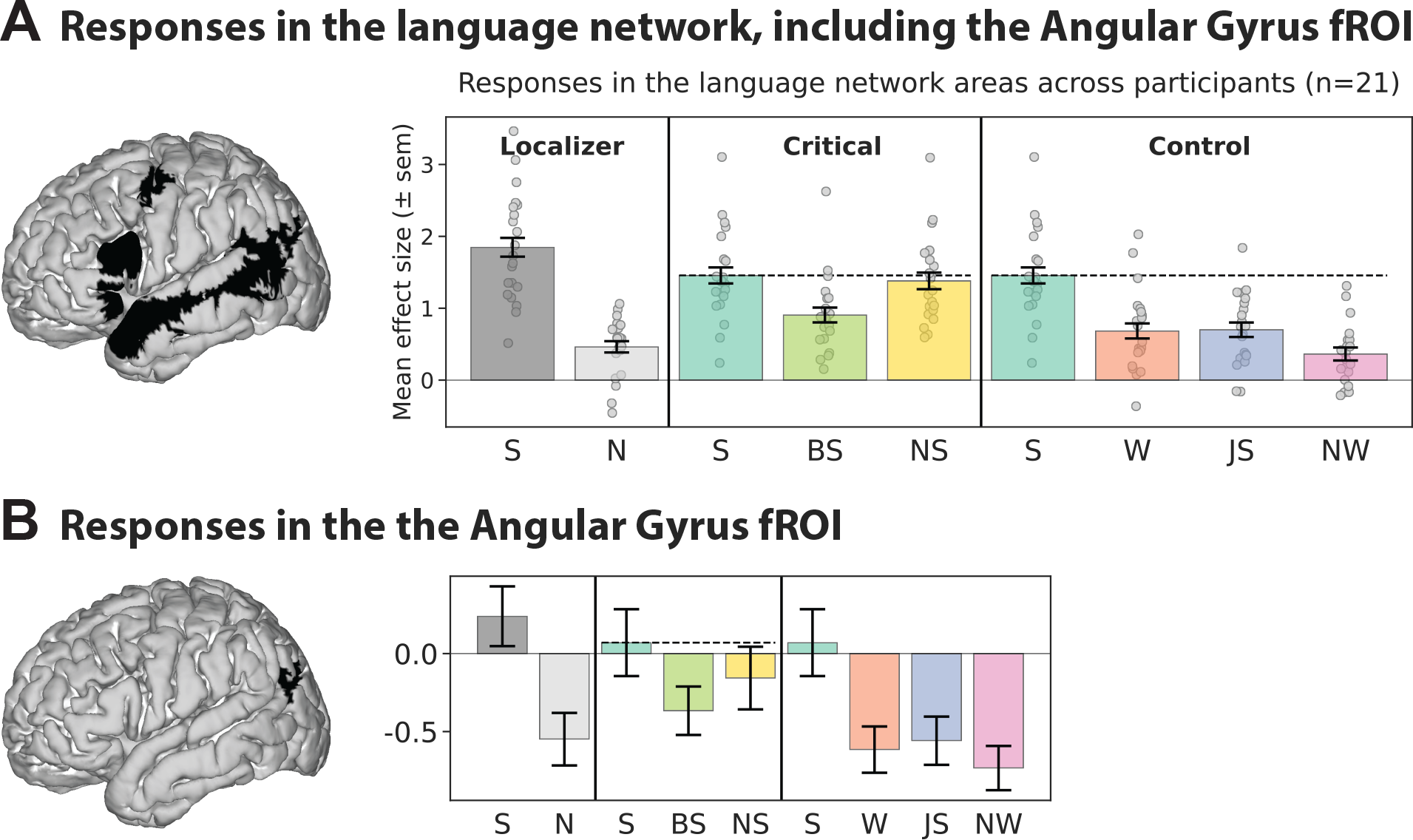
Responses in the language network, including the Angular Gyrus fROI. A) Neural responses (in % BOLD signal change relative to fixation) to the conditions of the language localizer and critical and control experimental conditions within the language network when including the Angular Gyrus fROI. **B)** Responses in just the Angular Gyrus fROI.

**Figure SI 7.**
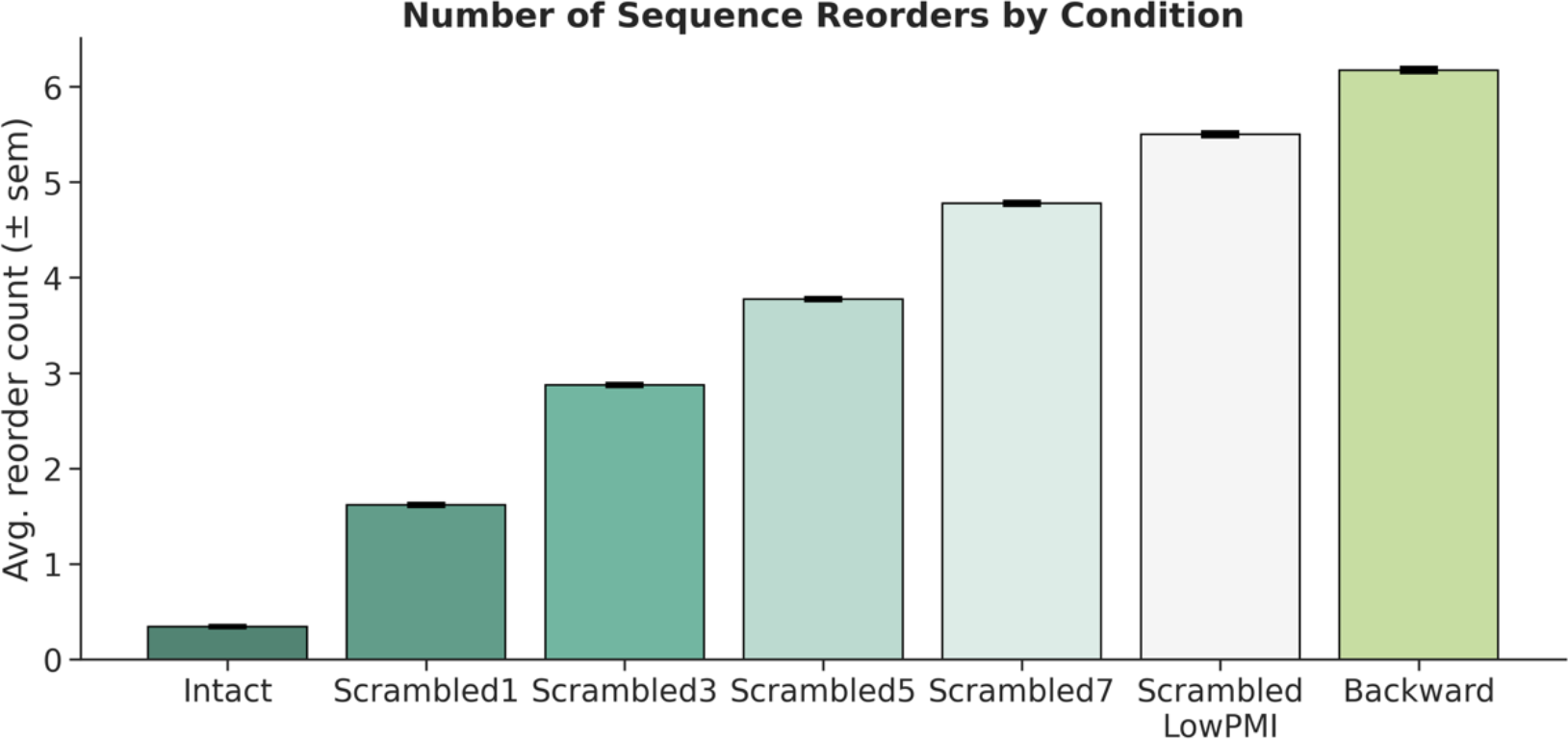
Validation of SynReco behavioral reconstruction paradigm. Participants actively reorder words during incremental sentence processing: each increase in the number of local word swaps led to an incremental increase in the number of time steps at which participants reordered the available words on the screen.

**Figure SI 8.**
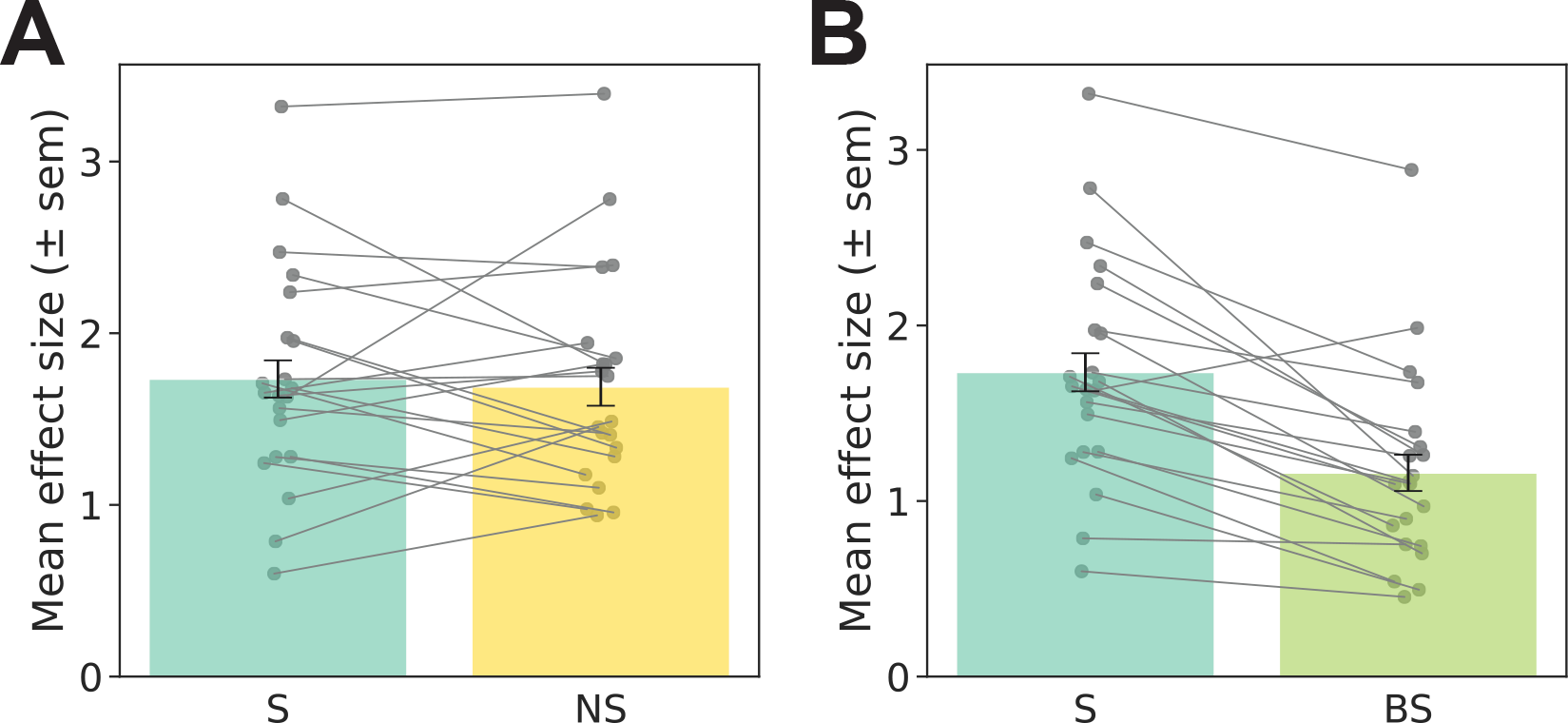
Individual subject effects for the critical conditions. Even though there is individual variability, the trends observed at the population level mostly hold in individual subjects, as well.

**Figure SI 9.**
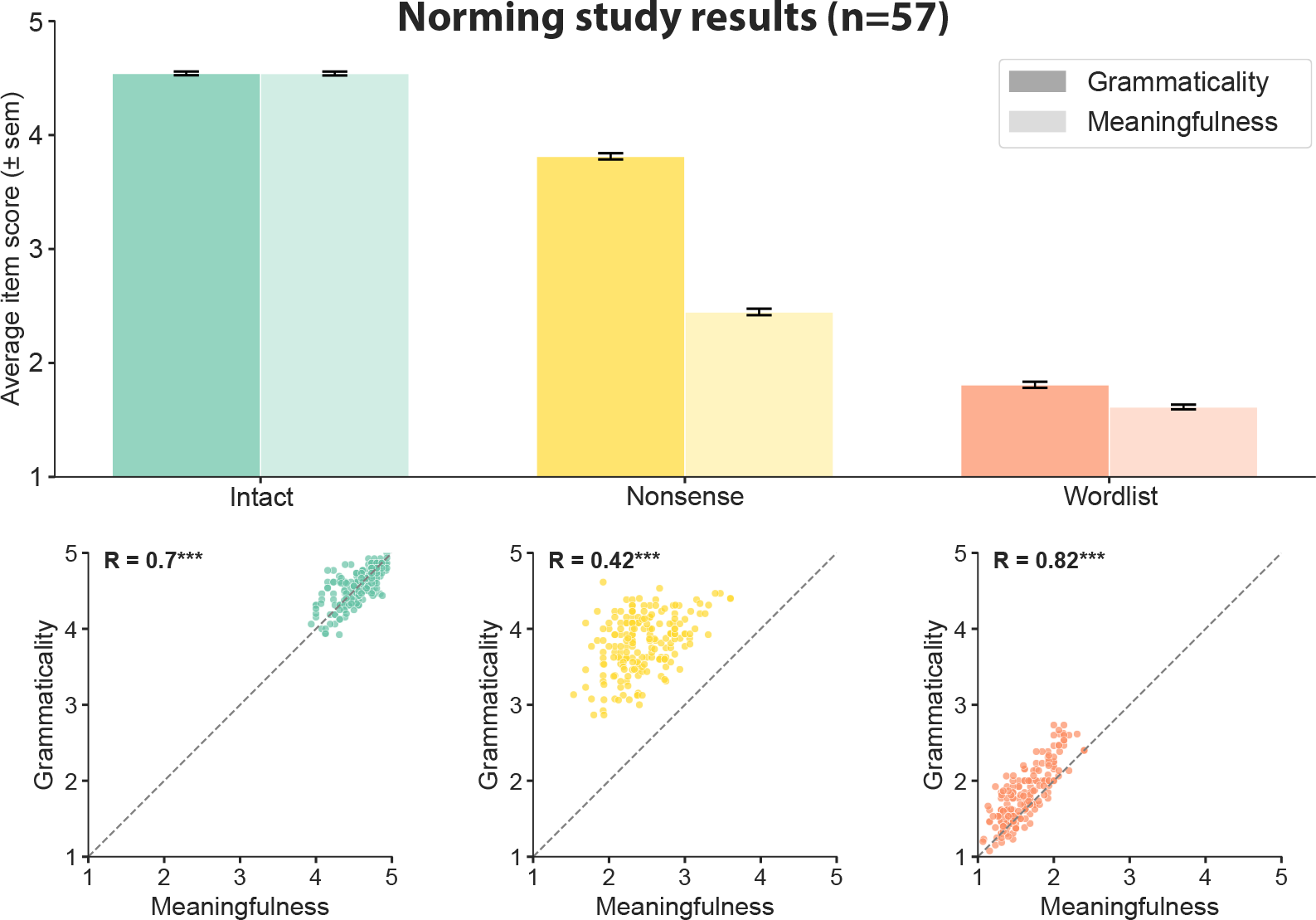
**Full set of results from the behavioral norming study (Figure 4A**). We asked participants to rate our *Plausible Sentence*, *Nonsense Sentence*, and *Word List* stimuli for two features: grammaticality and meaningfulness. *Nonsense Sentence* stimuli successfully dissociate the two features, even though they tend to be correlated. Nevertheless, *Nonsense Sentence* stimuli were rated worse grammatically than *Plausible Sentence* stimuli.

**Figure SI 10.**
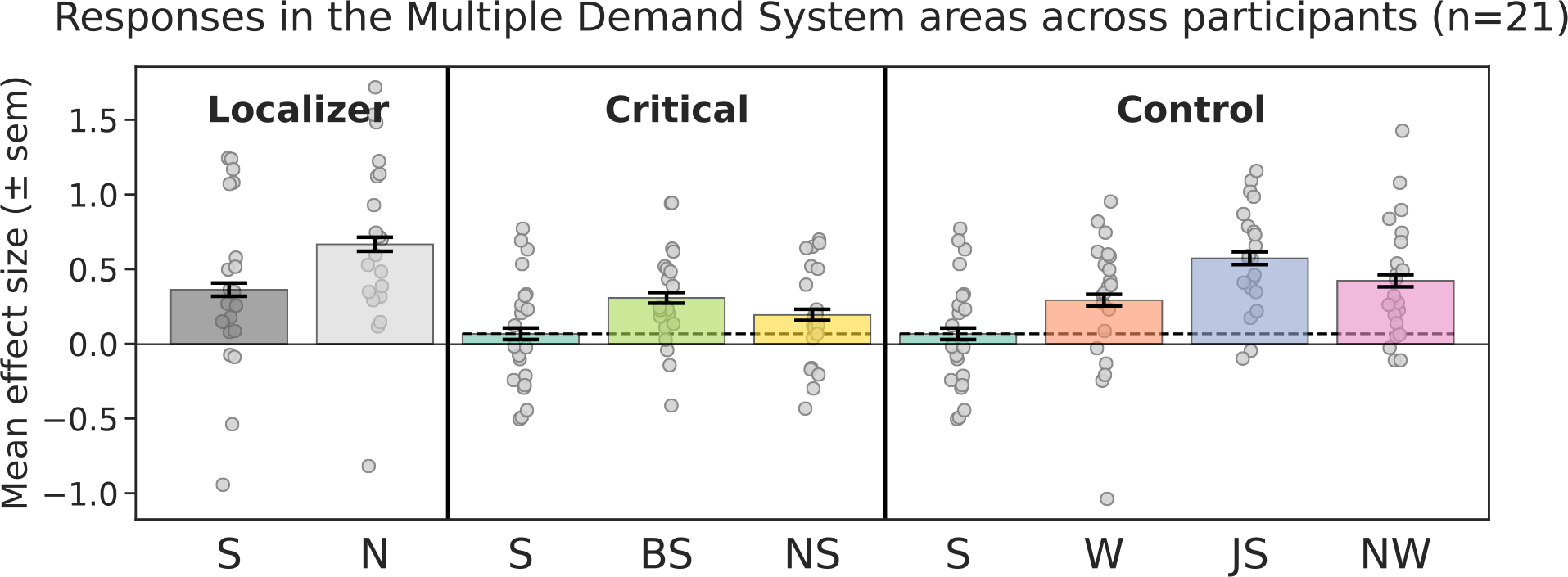
Multiple Demand system response. Neural responses (in % BOLD signal change relative to fixation) to the conditions of the language localizer and experimental conditions in the Multiple Demand (MD) system.

**Figure SI 11.**
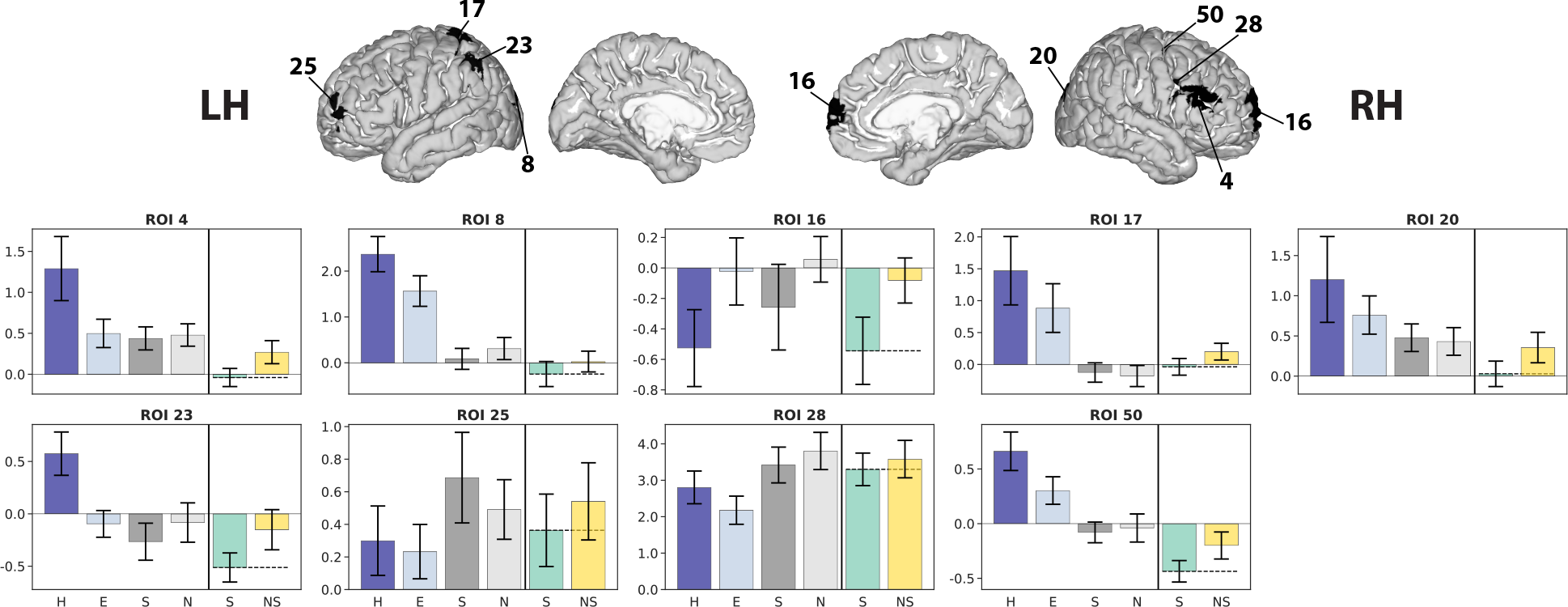
Finding regions that work harder when processing *Nonsense* sentences relative to *Plausible* sentences. The GSS whole-brain analysis for the *Nonsense sentence* > *Plausible sentence* contrast recovers a large network of brain regions. Follow-up analyses looking at the replicability of the contrast effect when including all subjects finds a subset of n=9 fROIs that show significant effects (shown here; average response shown in **Figure 4D**). We show the corresponding neural responses (in % BOLD signal change relative to fixation) within these fROIs to the conditions of the Multiple Demand (MD), language localizer and experimental conditions. However, these regions do not survive corrections for multiple comparisons across the entire network.

**Figure SI 12.**
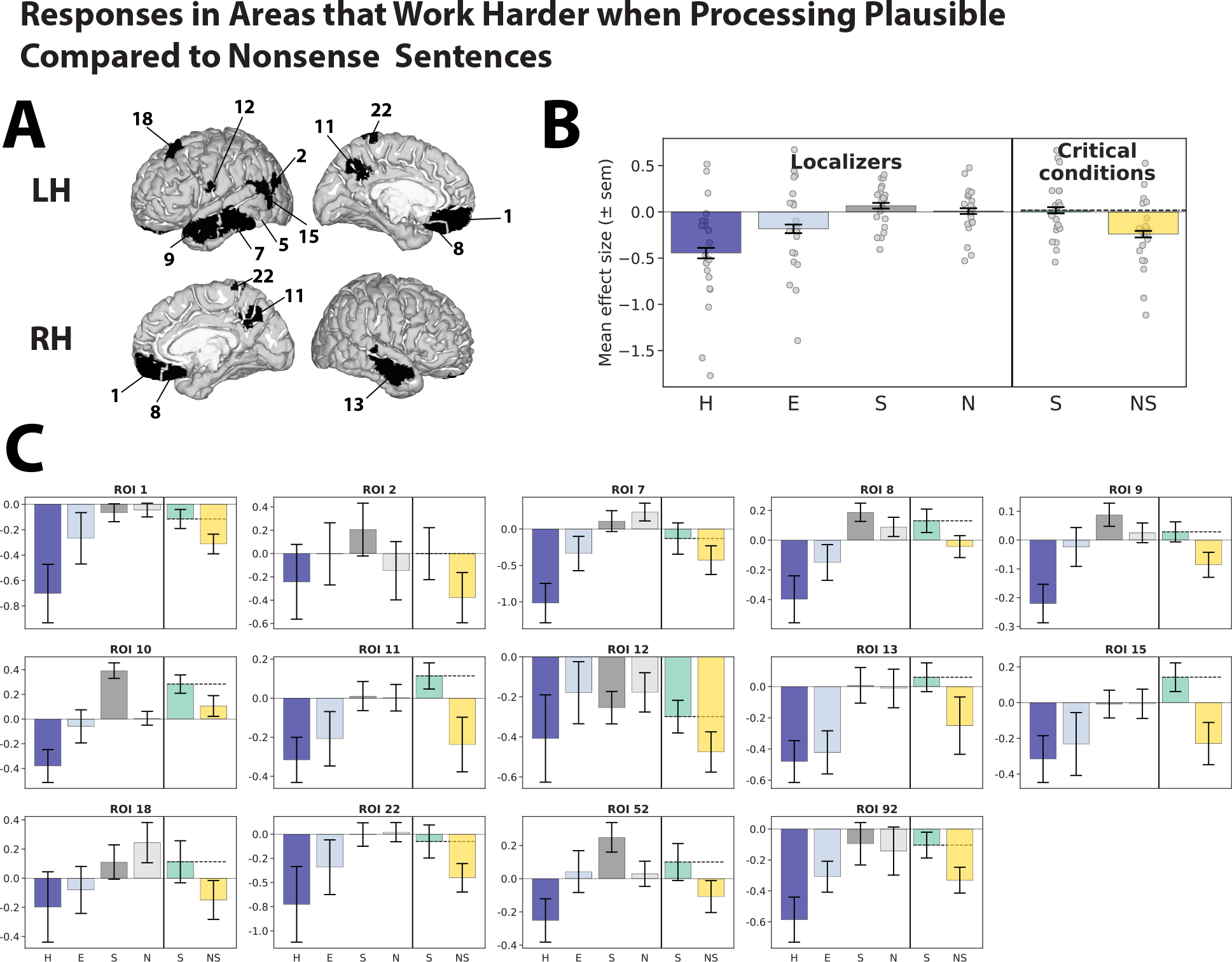
Finding regions that work harder when processing *Plausible* sentences relative to *Nonsense* sentences. A) The fROIs identified through GSS whole-brain analysis for the *Plausible sentence* > *Nonsense sentence* contrast, in which follow-up analyses looking at the replicability of the contrast effect when including all subjects show significant effects. However, only some of these regions survive corrections for multiple comparisons across the entire network. **B)** Neural responses (in % BOLD signal change relative to fixation) averaged across these fROIs to the conditions of the Multiple Demand (MD), language localizer and experimental conditions. **C)** Individual fROI responses.

**Figure SI 13.**
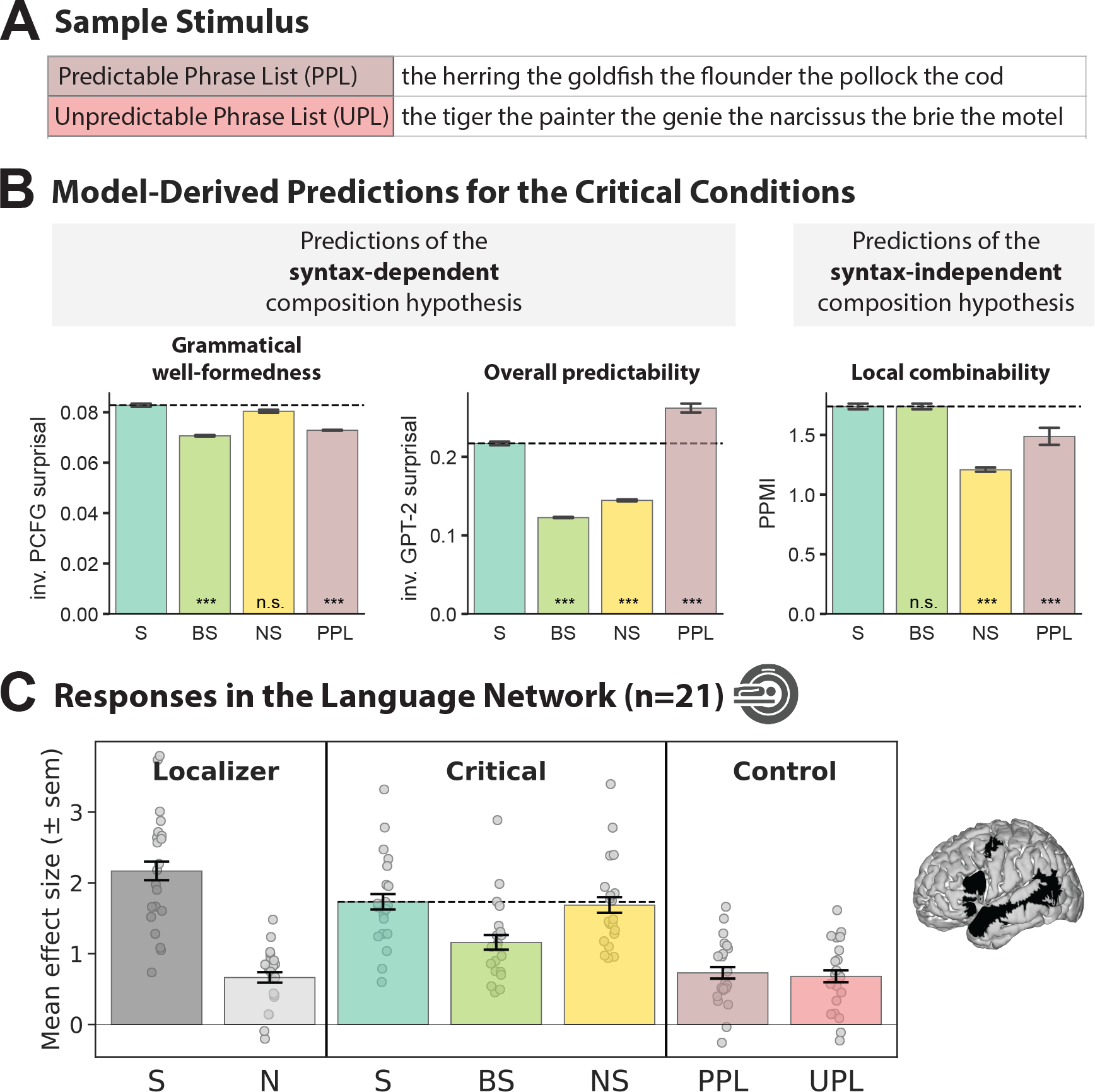
Language network response to semantically predictable and semantically unpredictable phrase lists. A) A sample item for the predictable and unpredictable phrase list conditions. Both conditions are made up of concatenated determiner phrases of the form *‘the noun’*; for the predictable phrase list condition (PPL), nouns are drawn from a semantically coherent category (here: kinds of fish), for control condition, unpredictable phrase lists (UPL), nouns are randomly drawn from the categories used to design the PPL stimuli. **B)** Quantitatively derived predictions for syntax-dependent vs. syntax-independent semantic composition hypotheses. The syntax-dependent panel is split up into predictions derived via structure-mediated vs. expectation-mediated incremental processing models (see <u>Discussion</u>). To match the expected direction of the neural responses in the language network, we show inverse surprisal (i.e., the reciprocal of surprisal) for the PCFG and GPT-2 models. Significant difference to the *Sentence* condition was established via post hoc pairwise t-tests, with p-values corrected for multiple comparisons using the Bonferroni procedure. **D)** Neural responses (in % BOLD signal change relative to fixation) to the conditions of the language localizer and critical and control experimental conditions within the language network (averaged across the five regions, see brain on the right). Dots show individual subject responses; error bars show standard errors of the mean by participants. The observed response shows that, in the absence of complex meaning, having a highly predictable stimulus (see panel B, Overall predictability) is insufficient to engage the language network.

### Supplementary Tables

**Table SI 1.** Statistics for PPMI stimuli characterization (<u>Results;</u>. **Section 3.1).** Pairwise, two-sided, dependent t-tests for all comparisons performed between the PPMI values for the *Intact* and all conditions of interest. P-values were corrected for multiple comparisons using the Bonferroni procedure. Effect sizes, as quantified by Cohen’s d are reported.

|  | Comparison with *Intact* |  |  |
| --- | --- | --- | --- |
|  | T-test statistic | Adjusted p-value | Cohen's d |
| Scr1 | 0.316 | 1.000 | 0.014 |
| Scr3 | 1.840 | 0.522 | 0.121 |
| Scr5 | 3.953** | 0.003 | 0.303 |
| Scr7 | 4.737*** | 0.000 | 0.575 |
| ScrLowPMI | 12.875*** | 0.000 | 2.207 |
| Backward | 0.201 | 1.000 | 0.000 |

**Table SI 2.**
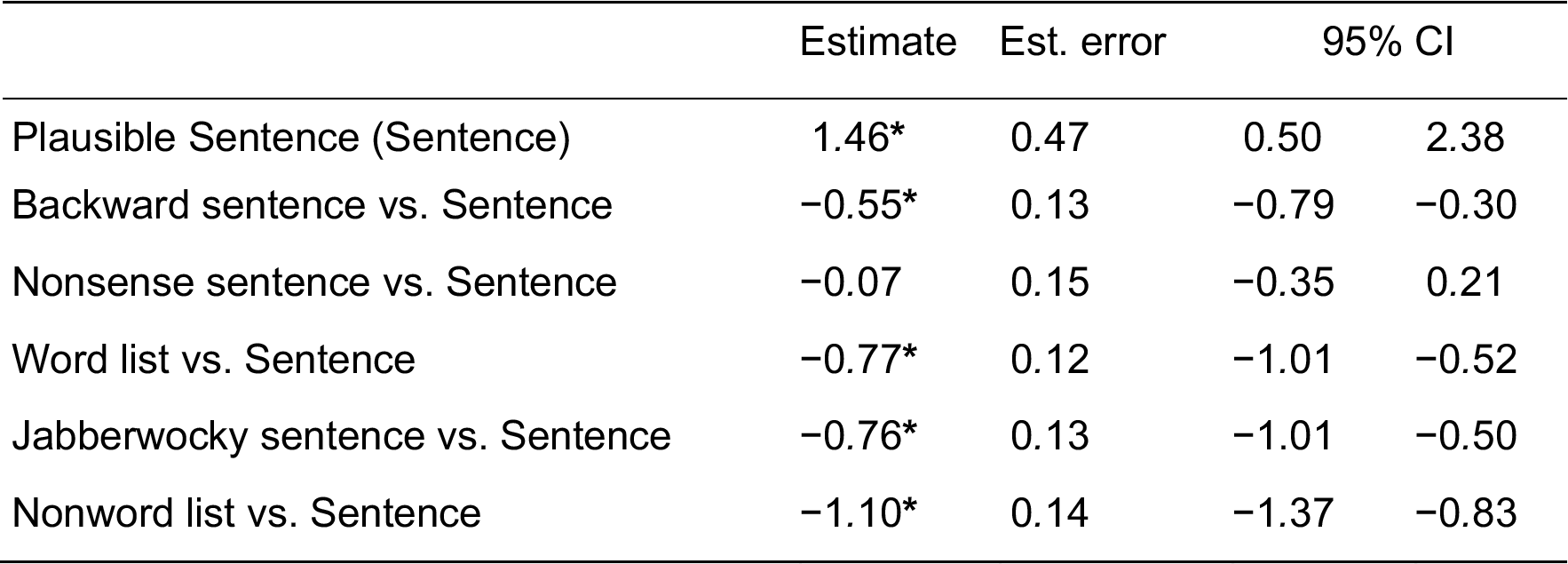
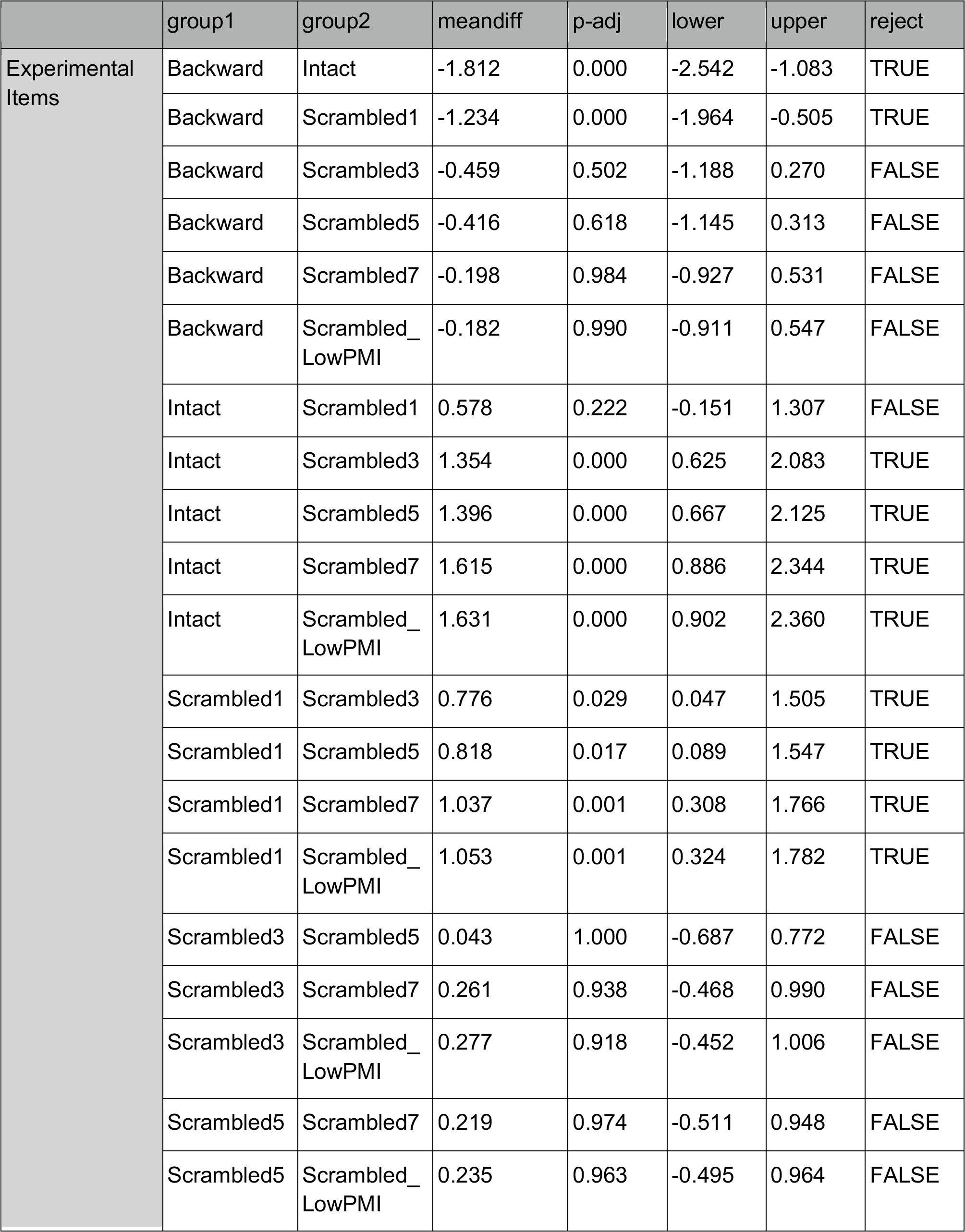
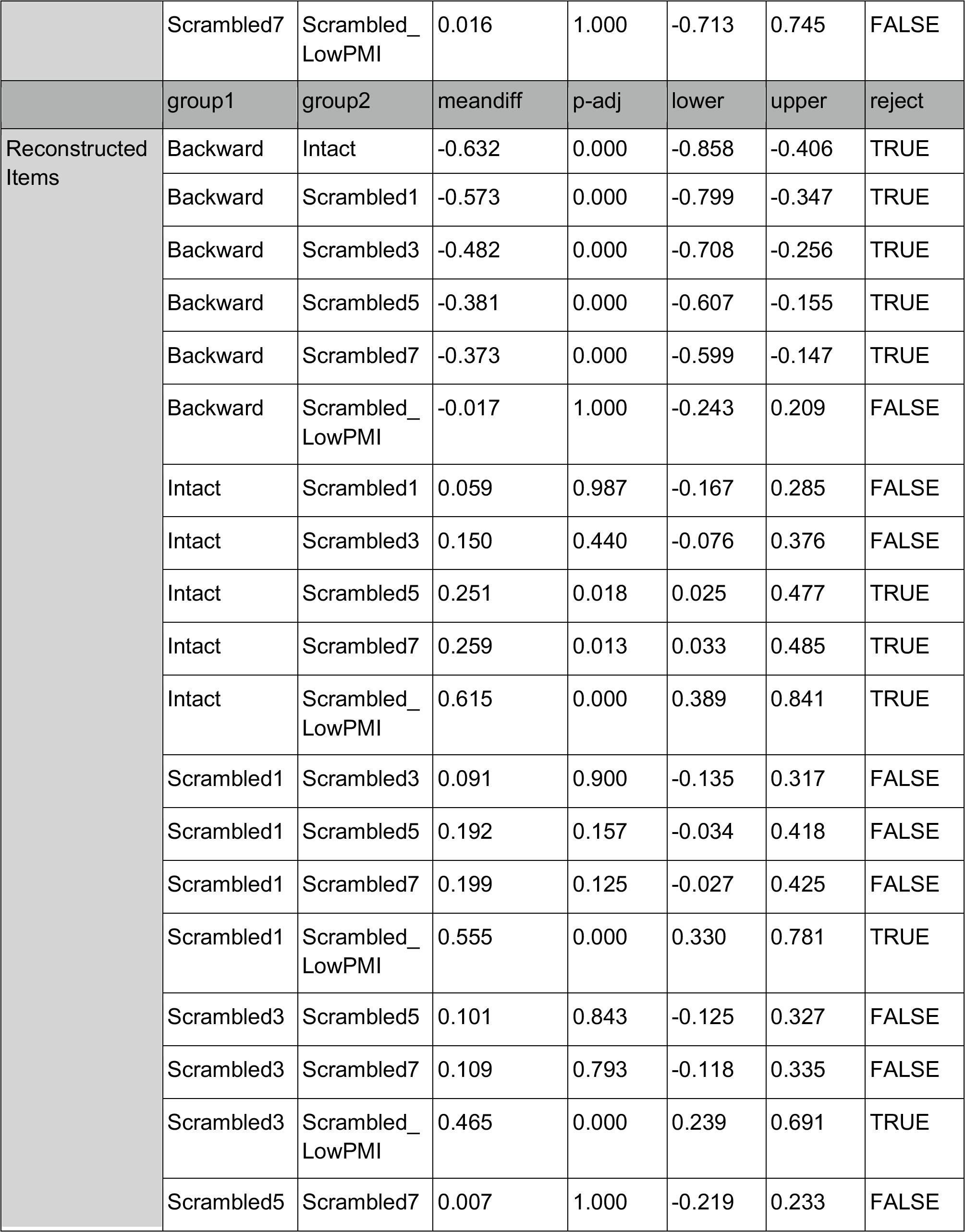
Results of mixed-effects linear regression for fMRI responses within the language network when including the Angular Gyrus fROI. Stimulus type was dummy-coded with *Sentence* as the reference level. **\***Denotes significant difference.

**Table SI 3.**
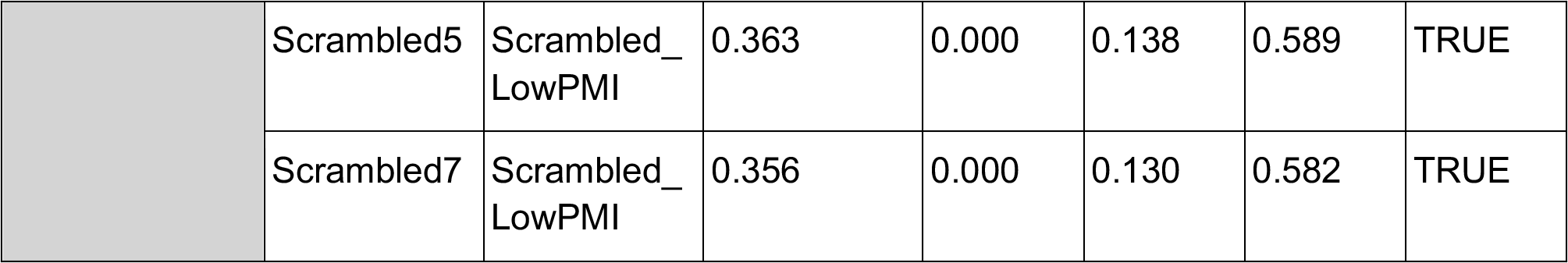
Statistics for. **Figure 1C in the main text (<u>Results;</u> Section 3.1).** Multiple comparisons of group means using Tukey’s Honestly Significant Difference (HSD) test.

**Table SI 4.** Statistics for verbatim reconstruction (<u>Results;</u>. **Section 3.1; Figure 1E).** The results of a mixed-effect logistic regression model with a fixed effect and random slopes for Condition, and random effects for Participant and Item. **\***Denotes significant difference.

|  | Estimate | Est. Error | 95% CI |  |
| --- | --- | --- | --- | --- |
| Grand mean | -0.74* | 0.13 | -1.00 | -0.47 |
| Scr1 - Int | -2.02* | 0.45 | -2.96 | -1.14 |
| Scr3 - Scr1 | -1.23* | 0.41 | -2.04 | -0.43 |
| Scr5 - Scr3 | -0.35* | 0.41 | -1.15 | 0.44 |
| Scr7 - Scr5 | -1.07* | 0.42 | -1.90 | -0.23 |
| ScrLowPMI - Scr7 | -1.46* | 0.46 | -2.38 | -0.57 |
| Backward<br>ScrLowPMI | -0.20 | 0.50 | -1.19 | 0.77 |

**Table SI 5.** The results of mixed effect linear regressions for the five language functional regions of interest (<u>Results;</u> Section 3.2). Condition was dummy-coded with *Sentence* as the reference level. IFGorb—orbital inferior frontal gyrus, MFG—middle frontal gyrus, AntTemp—anterior temporal lobe, PostTemp—posterior temporal lobe, Jab. — Jabberwocky.

|  |  | Estimate | Est.Error | 95% CI |  |
| --- | --- | --- | --- | --- | --- |
| IFGorb | Sentence (Sent) | 1.29* | 0.18 | 0.93 | 1.64 |
|  | Backward Sent vs. Sent | -0.69* | 0.12 | -0.93 | -0.45 |
|  | Nonsense Sent vs. Sent | -0.06 | 0.13 | -0.31 | 0.21 |
|  | Word List vs. Sent | -0.87* | 0.15 | -1.15 | -0.59 |
|  | Jab. Sent vs. Sent | -0.59* | 0.12 | -0.82 | -0.35 |
|  | Nonword List vs. Sent | -1.09* | 0.13 | -1.35 | -0.84 |
| IFG | Sentence (Sent) | 1.73* | 0.16 | 1.42 | 2.07 |
|  | Backward Sent vs. Sent | -0.72* | 0.13 | -0.97 | -0.46 |
|  | Nonsense Sent vs. Sent | 0.10 | 0.13 | -0.14 | 0.36 |
|  | Word List vs. Sent | -0.91* | 0.13 | -1.17 | -0.65 |
|  | Jab. Sent vs. Sent | -0.89* | 0.13 | -1.14 | -0.63 |
|  | Nonword List vs. Sent | -1.34* | 0.14 | -1.62 | -1.07 |
| MFG | Sentence (Sent) | 2.31* | 0.40 | 1.48 | 3.09 |
|  | Backward Sent vs. Sent | -0.27* | 0.14 | -0.55 | -0.02 |
|  | Nonsense Sent vs. Sent | -0.05 | 0.14 | -0.32 | 0.22 |
|  | Word List vs. Sent | -0.56* | 0.14 | -0.82 | -0.29 |
|  | Jab. Sent vs. Sent | -0.63* | 0.13 | -0.90 | -0.36 |
|  | Nonsense Sent vs. Sent | -0.05 | 0.14 | -0.32 | 0.22 |
| AntTemp | Sentence (Sent) | 1.20* | 0.10 | 1.02 | 1.40 |
|  | Backward Sent vs. Sent | -0.54* | 0.07 | -0.68 | -0.41 |
|  | Nonsense Sent vs. Sent | -0.11 | 0.08 | -0.26 | 0.04 |
|  | Word List vs. Sent | -0.73* | 0.07 | -0.86 | -0.59 |
|  | Jab. Sent vs. Sent | -0.77* | 0.08 | -0.92 | -0.62 |
|  | Nonword List vs. Sent | -1.04* | 0.10 | -1.22 | -0.84 |
| PostTemp | Sentence (Sent) | 2.12* | 0.16 | 1.80 | 2.42 |
|  | Backward Sent vs. Sent | -0.64* | 0.10 | -0.85 | -0.44 |
|  | Nonsense Sent vs. Sent | -0.12 | 0.11 | -0.33 | 0.10 |
|  | Word List vs. Sent | -0.90* | 0.10 | -1.10 | -0.69 |
|  | Jab. Sent vs. Sent | -1.03* | 0.11 | -1.24 | -0.81 |
|  | Nonword List vs. Sent | -1.37* | 0.11 | -1.59 | -1.15 |

**Table SI 6:** Results of mixed-effects linear regression for fMRI responses within the Multiple Demand network. Stimulus type was dummy-coded with *Sentence* as the reference level. Sent — Sentence, Jab. — Jabberwocky. **\***Denotes significant difference.

|  | Estimate | Est. error | 95% CI |  |
| --- | --- | --- | --- | --- |
| Sentence (Sent) | 0.07 | 0.13 | −0.19 | 0.31 |
| Backward Sent vs. Sent | 0.24* | 0.09 | 0.41 | 1.01 |
| Nonsense Sent vs. Sent | 0.12 | 0.09 | −0.04 | 0.29 |
| Word List vs. Sent | 0.22* | 0.11 | 0.00 | 0.45 |
| Jab. Sent vs. Sent | 0.50* | 0.09 | 0.33 | 0.67 |
| Nonword List vs. Sent | 0.36* | 0.10 | 0.15 | 0.57 |

## Supplementary Methods

### Behavioral incremental processing cost study

In this experiment, we measure the processing cost associated with the processing of grammatically well-formed sentences that convey plausible vs. unconventional meanings (i.e., the original sentences vs. the Nonsense versions of the sentences from the fMRI study).

### Paradigm, and design and materials

We used the Maze self-paced reading paradigm (Forster et al., 2009; Boyce et al., 2020). In this paradigm, stimuli are revealed one word at a time, and at each time step, the correct (target) word is accompanied by a contextually inappropriate distractor word, and participants have to indicate the word that they believe is a more likely continuation via pressing one of two buttons (**Figure 4B**). Boyce et al. (2020) showed that reaction times (RTs) in this paradigm effectively capture incremental processing cost.

The experiment included two conditions: the *Sentence* and *Nonsense Sentence* conditions from the fMRI study (192 stimuli per condition; for details see Section 2.2.1). The 384 stimuli were distributed across 4 experimental lists (96 stimuli each, 48 per condition) such that each list contained only one condition of an item. In addition to the critical stimuli, each list included 4 practice items. To generate the distractor words for each time step of each stimulus, we used the automatic implementation of Boyce et al. (2020), where the distractors are real words that are not grammatically licensed by the preceding content.

### Procedure

The experiment was implemented in the Maze module (Boyce et al., 2020) within the Ibex web-based psycholinguistic experiment software platform (https://github.com/addrummond/ibex).

The experiment began with detailed instructions. Following the instructions, participants completed 4 practice trials. Upon the completion of the practice trials, the critical experiment began. The 48 stimuli in each list were grouped into 6 blocks of 8 stimuli each, and participants were informed how many blocks remained after completing each block. To encourage participants to stay attentive throughout the experiment, a delay period of 2 s prevented participants’ keypresses from registering whenever an error was made (for motivation, see https://vboyce.github.io/Maze/delay.html). The average completion time was ∼10.5 min.

### Participants

We recruited 80 participants through the Prolific web-based testing platform, restricting our task to participants with IP addresses in the United States. Participants were excluded from the analyses if their performance on the task was low (<80% accuracy; average accuracy was >90%). Data from 70 participants were included in the final analysis.

